# Molecular Insights Into PGPR-Biochar–Mediated Arsenic Detoxification and Nutritional Enhancement in Rice Rhizospheres

**DOI:** 10.1101/2025.08.11.669651

**Authors:** Arnab Majumdar, Tarit Roychowdhury, Ioly Kotta-Loizou, Martin Buck

## Abstract

Arsenic (As) contamination in paddy soils threatens rice productivity and food safety through enhanced As(III) mobility under anaerobic conditions. This study evaluated the synergistic effects of rice biochar (RBC) and plant growth-promoting rhizobacteria (PGPR) on arsenic bioavailability and rice physiological responses under controlled and field conditions. Rice straw biochar pyrolyzed at 400 °C exhibited high porosity and surface functionality, while Metabacillus indicus AMB4 demonstrated arsenic tolerance (≤3500 mg kg^-1^) and multiple PGPR traits (IAA: 45.2 μg mL^-1^, phosphate solubilization index: 2.8, ACC deaminase: 18.7 nmol α-ketobutyrate mg^-1^ h^-1^). Combined RBC+PGPR treatment reduced soil arsenic bioavailability by 85% (p<0.001) and plant arsenic accumulation by 78% compared to controls, while significantly enhancing antioxidant enzyme activities (CAT: 4.2-fold, SOD: 3.7-fold, APX: 3.1-fold increase). Metagenomics revealed restructured microbial communities with increased Proteobacteria abundance and upregulated arsenic detoxification genes (arsC, arsB, aioA) coupled with enhanced nutrient cycling pathways (nifH, phoD, soxA). Network analysis demonstrated functional integration between As-transformation and nutrient metabolism, with coordinated expression of 47 arsenic-responsive and 126 nutrient-cycling genes. Field trials confirmed superior plant performance with RBC+PGPR treatment yielding 56% more tillers and 73% more panicles than controls, while maintaining enhanced soil nutrient status. This integrated biochar-PGPR system provides a mechanistically robust approach for simultaneous arsenic remediation and crop productivity enhancement in contaminated rice ecosystems, demonstrating practical applicability for sustainable paddy agriculture.

## Introduction

Arsenic (As) contamination in paddy soils constitutes a major constraint to rice (Oryza sativa) productivity and food safety, primarily due to the high mobility of arsenite [As(III)] under anaerobic flooded conditions typical of paddy fields. Among advanced biogeochemical remediation strategies, the combined application of plant growth-promoting rhizobacteria (PGPR) and straw–derived rice biochar (RBC) has garnered considerable attention for its efficacy in reducing As bioavailability while simultaneously enhancing nutrient cycling within the rhizosphere. Rice straw biochar, particularly pyrolyzed at 400–500□°C, exhibits a highly porous structure and substantial surface functional groups that facilitate strong adsorption of arsenic species, including As(III) and As(V), through mechanisms such as surface complexation, ion exchange, and co-precipitation with iron and silica oxides forming in situ (Chuan et al., 2017). The biochar-induced increase in pH and modulation of redox potential (Eh) transform rhizosphere As speciation, promoting the formation of iron plaques on rice root surfaces that act as reactive barriers sequestering As and restricting its translocation to aerial tissues (Chen et al., 2021). Functionally, RBC fosters enhanced microbial oxidation of As(III) via activation of the aioA gene in As-oxidizing bacteria, contributing to conversion of more toxic arsenite to less mobile arsenate (Li et al., 2020). Moreover, RBC’s porous matrix and labile carbon content provide habitat and nutrition that substantially increase soil microbial biomass, fostering a diverse community including Fe(III)- and As(V)-reducing bacteria such as Geobacter and Shewanella, which influence electron transfer and As biogeochemical cycling under anaerobic soil conditions (Sun et al., 2024).

Colonization of RBC-amended soils by PGPR enriches expression of As-transforming genes, including arsC (arsenate reductase), arrA (respiratory arsenate reductase), aioA (arsenite oxidase), and arsM (arsenic methyltransferase), enabling multifaceted As detoxification mechanisms such as arsenate reduction, arsenite oxidation, and volatilization of methylated products like dimethylarsine and trimethylarsine, which diffuse away from the rhizosphere, effectively reducing soil As load (Mukhopadhyay et al., 2021). In addition to detoxification, PGPR perform pivotal nutrient cycling functions—nitrogen fixation via nifH, phosphate solubilization via phoD, siderophore production for iron acquisition, auxin (IAA) synthesis for root growth stimulation, and ACC deaminase activity to mitigate ethylene stress—thus promoting plant growth under arsenic stress (Zhou et al., 2022). High-throughput sequencing and network analyses reveal that PGPR and RBC synergistically restructure the rice rhizosphere microbiome, establishing functional gene clusters linking As detoxification pathways with nutrient cycling that suggest co-selection and stable functional integration under biochar amendment.

This integrated PGPR–RBC strategy, by simultaneously modulating geochemical conditions, microbial community structure, functional gene expression, and plant physiology, provides a rigorous, multi-mechanistic approach for mitigating As toxicity and enhancing nutrient acquisition in rice paddy ecosystems, representing a promising sustainable technology for arsenic-contaminated soil remediation and food safety assurance. Based on the reports of biochar application and PGPR involvement in plant growth, three knowledge gaps were identified, and the objectives of this study were defined accordingly: (i) How to manage the rice straw generated after its harvest(ii) How can biochar influence the plant’s anatomy while conferring induction on the field soil microbiome(iii) Can biochar-doped PGPR increase As detoxification and modulate field soil nutrientsThese points will help to underpin the practical applicability of biochar combined with a PGPR as an optimum mode of As mitigation and nutrient enrichment in natural environment.

## Materials and Methods

### Biochar preparation and characterisation

Dry rice shoots obtained from local farmers were finely chopped and oven-dried using a hot air system. Pyrolysis was subsequently carried out under limited oxygen conditions, with the temperature increased at a rate of 20□°C□min□¹ until reaching 400□°C, which was maintained for 45□minutes (method adapted from Naeem et□al., 2017). Following pyrolysis, the resulting rice biochar (RBC) was cooled to ambient temperature and sieved through a 2□mm mesh to ensure homogeneity. The arsenic (As) adsorption capacity of the RBC was then evaluated using energy-dispersive X-ray spectroscopy (EDX; Oxford INCA) coupled with field emission scanning electron microscopy (FE-SEM; Carl Zeiss SUPRA 55VP). EDX analysis was performed on three sample types: untreated RBC (control), As-adsorbed RBC, and As-adsorbed RBC inoculated with plant growth-promoting rhizobacteria (PGPR), both for qualitative visualisation and quantitative elemental assessment.

### PGPR screening and As tolerance analysis

Fresh soil samples were collected 5–10 cm below the surface and stored at 4°C until processing. One gram of soil was added to 9 mL of sterile distilled water to create a 10□¹ dilution, which was vortexed for 2–5 minutes to release bacteria from soil particles. Serial ten-fold dilutions were prepared up to 10□□ or 10□□ by transferring 1 mL into fresh sterile diluent. From each dilution, 100 μL was spread onto nutrient agar plates using sterile spreaders, and plates were incubated inverted at 30–37°C for 24–48 hours. Well-isolated bacterial colonies were selected from higher dilution plates and purified through repeated streak plating until single colonies were obtained. Pure cultures were preserved on agar slants or in glycerol broth at −20°C for long-term storage, with further characterization performed. Fresh bacterial colonies were harvested from pure cultures and subjected to genomic DNA extraction using a commercial kit or standard phenol-chloroform method. The 16S rRNA gene was amplified by polymerase chain reaction (PCR) using universal bacterial primers targeting conserved regions (e.g., 27F: 5’-AGAGTTTGATCMTGGCTCAG-3’ and 1492R: 5’-TACGGYTACCTTGTTACGACTT-3’). Bacterial identification was assigned based on sequence similarity thresholds, with ≥98% similarity. Two isolated colonies were selected for this identification-AMB2 and AMB4.

Arsenic Tolerance and Cellular Bioaccumulation-Strains AMB2 and AMB4 were exposed to concentration gradients of sodium arsenite (NaAsO□) at 100, 250, 400, 600, 800, 900, 1000, 1100, and 1200 mg kg□¹, and sodium arsenate dibasic heptahydrate (Na□HAsO□·7H□O) at 100, 400, 800, 1000, 1500, 2000, 2500, 2800, 3000, and 3500 mg kg□¹ on NA plates. Growth responses were assessed after 24 and 48 hours of incubation. Intracellular arsenic accumulation was quantified using a modified acid digestion protocol (Banerjee et al., 2013). Bacterial cultures were grown in NB medium supplemented with arsenate (10-30 mg L□¹) or arsenite (4-10 mg L□¹) for 72 hours at 35°C. Daily cell harvests were obtained by centrifugation, and the resulting pellets were acid-digested using 65% nitric acid (HNO□) until complete dissolution. Digested samples were diluted to 10 mL with Milli-Q water and analyzed for total intracellular arsenic content using inductively coupled plasma mass spectrometry (ICP-MS, Thermo Scientific Q-ICP-MS, XSeries 2). After observing the results of this As tolerance test, AMB4 was selected to be used in the experiments further.

### Assessment of PGPR traits of the selected AMB4

AMB4 isolate was evaluated for multiple beneficial traits using established protocols. Indole-3-acetic acid (IAA) production was quantified using the Salkowski reagent method, where bacterial cultures were grown in nutrient broth supplemented with 2 g L□¹ L-tryptophan at 28°C for 72 hours under dark conditions with shaking at 150 rpm (Singh et al., 2019; Jha et al., 2020). Cell-free supernatants (1 mL) were mixed with equal volumes of Salkowski reagent (0.5 M FeCl□ in 35% HClO□) and incubated for 30 minutes in darkness, with IAA concentrations determined spectrophotometrically at 530 nm against standard IAA curves (Singh et al., 2019; Ahmed et al., 2021). Gibberellin (GA) production was assessed by cultivating bacterial isolates in sterile nutrient broth for 10 days at 30°C with shaking at 100 rpm, followed by spectrophotometric analysis of cell-free supernatants using established colorimetric methods for GA quantification (Sagar et al., 2018; Nett et al., 2016). Siderophore production was evaluated using the chrome azurol sulfonate (CAS) assay, where bacterial supernatants (100 μL) were mixed with CAS reagent (100 μL) in 96-well microplates and incubated for 20 minutes before measuring absorbance at 630 nm, with siderophore units calculated as percent siderophore units (PSU) = [(Ar - As)/Ar] × 100, where Ar and As represent absorbance of reference and sample, respectively (Louden et al., 2011; Machuca & Milagres, 2003; Pérez-Miranda et al., 2007). Phosphate solubilization was determined both qualitatively and quantitatively using National Botanical Research Institute’s phosphate (NBRIP) medium containing tricalcium phosphate as the sole phosphorus source, with solubilization index calculated as (colony diameter + halo diameter)/colony diameter, and quantitative phosphate release measured colorimetrically using the vanadate-molybdate method at 430 nm (Nautiyal, 1999; Kumar et al., 2023; Vikram et al., 2007). ACC deaminase (ACCD) activity was assessed by growing bacterial cultures on Dworkin-Foster minimal medium containing 3 mM 1-aminocyclopropane-1-carboxylate (ACC) as the sole nitrogen source, with enzyme activity quantified through α-ketobutyrate production using the 2,4-dinitrophenylhydrazine assay and measuring absorbance at 540 nm against standard α-ketobutyrate curves (Belimov et al., 2005; Penrose & Glick, 2003; Siddikee et al., 2010). All assays were performed in triplicate with appropriate positive and negative controls to ensure reproducibility and accuracy of results.

### Total and bioavailable elemental analysis in soil-plant systems

Total elemental composition of soil samples was determined using X-ray fluorescence (XRF) spectroscopy (Bruker S8 Tiger, Germany; DST-FIST facility, IISER Kolkata). Plant samples were subjected to di-acid digestion using nitric acid and hydrogen peroxide (Marin et al., 2011; Majumdar et al., 2018) followed by analysis using inductively coupled plasma mass spectrometry (ICP-MS, Thermo Scientific X Series 2). For accurate quantification, certified reference materials were employed for instrument calibration. Four standards (JSD-1, SDC-1, MESS-3, and NIST 2711a) were used for major element calibration, while NIST 2711a and MESS-3 were utilized for trace element analysis. All samples and standards were oven-dried at 60°C for 48 hours and ground to fine powder using an agate mortar and pestle. Soil samples were sieved through 200 ASTM mesh, after which 4 g of sample was thoroughly mixed with 1 g of boric acid powder to ensure complete homogenization and eliminate clumping that could cause background scattering interference. The homogenized mixture was compressed into pellets using a hydraulic press, and XRF analysis was conducted with the smooth pellet surface oriented toward the X-ray beam path for optimal signal detection. Sequential extraction of soil-bound elemental fractions was performed following the modified Tessier protocol (Tessier et al., 1979; Sarkar et al., 2017). This procedure utilized a series of extractants including magnesium chloride, sodium acetate, acetic acid, ammonium hydroxide–hydrochloric acid, nitric acid, hydrogen peroxide, ammonium acetate, hydrofluoric acid, and perchloric acid to isolate five operationally defined fractions: exchangeable, carbonate-bound, Fe–Mn oxide-bound, organic matter-bound, and residual. The exchangeable and carbonate-bound fractions were considered bioavailable due to their ready mobilization under conditions of ionic exchange or pH/redox variations.

### Electron microscopy and ultrastructure observation of rice plants

Fresh plant shoot samples were sectioned into thin cross-sections using a microtome and subsequently air-dried under sunlight to remove excess intercellular moisture. Following complete dehydration, samples were sputter-coated with gold using an Edwards S150B sputter coater under vacuum conditions with pure argon gas. Gold coating was applied at 2000 V and 15 mA for 10 seconds, resulting in a uniform 2 nm thick conductive layer. Coated specimens were examined using field emission scanning electron microscopy (FE-SEM, Zeiss SUPRA 55VP) at controlled magnifications, with imaging focused specifically on the vascular architecture of the plant tissues (Majumdar et al., 2019).

### Metagenomics and high-throughput sequencing of soil samples

Soil samples were taken directly from the study locations and rapidly chilled on dry ice to maintain molecular integrity until processing. Genomic DNA was extracted from each sample using the DNeasy PowerSoil Kit (Qiagen) in accordance with the manufacturer’s recommendations. DNA eluates were obtained in nuclease-free water, and sample concentration and purity were evaluated using a NanoDrop Lite spectrophotometer (Thermo Scientific). Only samples meeting quality thresholds were advanced to library construction. For this step, Oxford Nanopore Technologies’ Ligation Sequencing Kit (SQK-LSK109) and PCR Barcoding Kit (EXP-PCR096) were applied. Amplicons underwent end-repair and dA-tailing via the NEBNext Ultra II End Repair/dA-Tailing Module (New England Biolabs, MA), followed by purification with AMPure XP magnetic beads (Beckman Coulter, USA). Barcoded libraries were combined and subjected to an additional end-repair cycle before a final cleanup using the NEB Blunt/TA Ligase Master Mix protocol. Sequencing runs were performed on the GridION X5 platform with R9.4.1 SpotON flow cells (FLO-MIN106), generating raw FAST5 files. Base-calling and demultiplexing were carried out using Guppy v2.3.4. The resulting sequence data were processed to identify operational taxonomic units, assess microbial community diversity, and annotate functional genes and pathways through the Kyoto Encyclopedia of Genes and Genomes (KEGG). Additional analyses included arsenic-responsive gene network mapping and nitrogen-phosphorus-sulfur transporter gene expression profiling.

### Statistical justifications and result analysis

All soil measurements were carried out in triplicate, and differences among treatments were evaluated for significance at a threshold of p ≤ 0.05. To examine overall patterns in the data, a one-way analysis of variance (ANOVA) was applied. In addition, the Tukey post-hoc test was performed across the complete dataset. For a more conservative interpretation of statistical significance, the Bonferroni correction was employed as a post-hoc adjustment to the soil–plant dataset. Given the inclusion of 4 experimental sites, 4 variables were considered in the correction, yielding an adjusted α-value of p ≤ 0.012. Microbial interaction networks were constructed in Cytoscape using the STRING database, which also enabled visualization of associated protein structures via the Protein Data Bank (PDB). To explore functional relationships, gene ontology (GO) terms from the experimental dataset were integrated with online resources through ShinyGO, allowing identification of connectivity among arsenic-related physiological responses and their protein products. Statistical analysis, data visualization, microbial diversity profiling, GO term mapping, and KEGG pathway analysis were conducted using a combination of software platforms, including GraphPad Prism 6, OriginPro 2024, SigmaPlot v12, ClustVis v2.0 (online), ShinyGO v0.77 (online), and Cytoscape v3.8.2.

## Results and Discussion

### Doping of PGPR with RBC and As adsorption

Figure 1 presents high-resolution Field Emission Scanning Electron Microscopy (FE-SEM) images alongside Energy Dispersive X-ray (EDX) spectroscopic analysis, illustrating the morphological and elemental changes in rice biochar (RBC) and plant growth-promoting rhizobacteria (PGPR)-doped RBC during arsenic adsorption. The FE-SEM micrograph of pristine RBC (Fig. 1a) reveals a highly porous and heterogeneous surface morphology characterized by an interconnected network of macropores and micropores, typical for rice straw-derived biochar subjected to pyrolysis at moderate temperatures (400–500°C) [Chen et al., 2021; Sun et al., 2024]. This porous structure offers an extensive surface area conducive to adsorptive interactions with contaminants. The accompanying EDX spectrum confirms the primary elemental composition dominated by carbon (C) and oxygen (O), with minor contributions of elements such as potassium (K), silicon (Si), aluminum (Al), and calcium (Ca), originating from inherent mineral components of biochar feedstock [Wang et al., 2020]. The absence of arsenic peaks indicates the uncontaminated surface baseline. After exposure to arsenic-laden solutions, the RBC surface undergoes discernible morphological changes (Fig. 1b). The biochar pores and surface irregularities appear partially occluded by fine precipitates or adsorbed mineral layers, consistent with previous observations of arsenic immobilization through surface complexation and co-precipitation [Li et al., 2020; Chuan et al., 2017]. EDX spectra here distinctly reveal the presence of arsenic (As) peaks, confirming successful adsorption onto RBC. Notably, increases in Fe and Si peak intensities suggest formation of iron-silica complexes and iron plaques, which are known to play a significant role in arsenic sequestration by biochar in paddy soils [Chen et al., 2021; Sun et al., 2024]. The physical sorption and chemical binding to these mineral surfaces reduce arsenic bioavailability and mobility.

**Figure 1.**
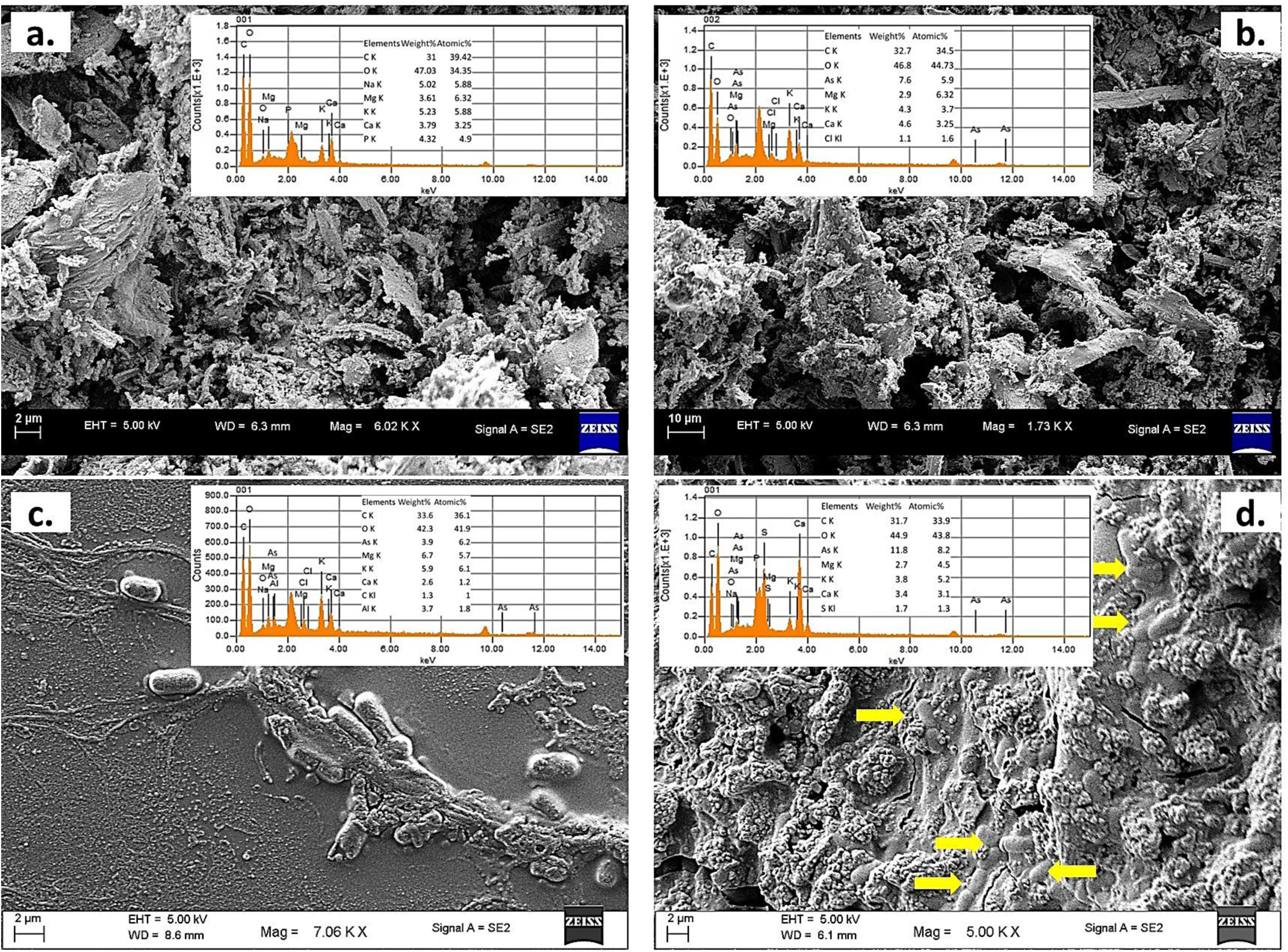
FE-SEM observation with EDX analysis of the RBC and PGPR-doped RBC As adsorption removal. Figure 1a-d sequentially represents control RBC, As-sorbed RBC, As-sorbed PGPR and As-sorbed RBC+PGPR.

In the micrographs of PGPR-treated surfaces without biochar (Fig. 1c), rod-shaped bacterial cells are visible forming biofilms and extracellular polymeric substances around cellular aggregates. This microbial colonization increases surface roughness and heterogeneity, enhancing sorption sites for arsenic through binding to microbial cell walls rich in functional groups such as carboxyl, phosphoryl, and hydroxyl [Mukhopadhyay et al., 2021; Zhou et al., 2022]. The EDX reveals abundant oxygen and carbon peaks consistent with organic biomass, while arsenic peaks indicate that microbial metabolism and cell surface adsorption contribute to arsenic immobilization. PGPR also express genes (e.g., arsC, aioA) enabling arsenic transformation and potentially volatilization, providing biotransformation pathways complementary to mere physical adsorption [Mukhopadhyay et al., 2021]. The RBC biochar doped with PGPR (Fig. 1d) exhibits a complex surface morphology where biochar pores are colonized by dense microbial aggregates, evidenced by rod-shaped bacterial cells embedded within a porous biochar matrix. Yellow arrows highlight these bacterial colonization sites, biofilm matrices, and mineral precipitates, implying a symbiotic structuring effect. The presence of PGPR on RBC surfaces likely enhances arsenic removal through combined physicochemical and biological pathways: biochar provides adsorption sites and habitat, while PGPR catalyze arsenic oxidation (aioA gene) and methylation (arsM gene), facilitating transformation of As(III) to less toxic or volatile forms [Li et al., 2020; Mukhopadhyay et al., 2021]. EDX analysis indicates stronger arsenic signals, confirming higher arsenic accumulation compared to RBC or PGPR alone, as well as elevated Fe and Si contents suggesting intensified formation of iron plaques that further immobilize arsenic [Chen et al., 2021]. Enhanced potassium and calcium peaks point to improved nutrient availability linked to PGPR bioactivity [Zhou et al., 2022]. The FE-SEM and EDX findings corroborate that rice-straw biochar provides a robust adsorptive substrate with extensive porosity and mineral phases favorable for arsenic sequestration. PGPR inoculation modifies this system, not only by biofilm-related adsorption but also via enzymatic arsenic transformations that reduce toxicity and mobilize volatile derivatives. The combined RBC+PGPR system creates a multifunctional microenvironment integrating physical sorption, microbial biotransformation, and nutrient cycling, leading to a synergistic effect on arsenic detoxification in contaminated paddy soils. This supports the concept that biochar amendments coupled with rhizobacterial consortia offer sustainable strategies for mitigating arsenic risks in rice agroecosystems [Li et al., 2020; Mukhopadhyay et al., 2021; Zhou et al., 2022].

### Antioxidant biochemical responses in pot and field-grown plants

The presented data (Figure 2a–l) detail the enzymatic and biochemical responses of rice plants subjected to arsenic (As) stress and treated with rice straw biochar (RBC) and PGPR, both in pot (a–f) and field (g–l) experiments. The measured parameters—MDA (malondialdehyde), protein content, and antioxidant enzymes CAT (catalase), SOD (superoxide dismutase), APX (ascorbate peroxidase), and GPX (guaiacol peroxidase)—serve as biomarkers for oxidative stress and detoxification activity in plant tissues.

**Figure 2.**
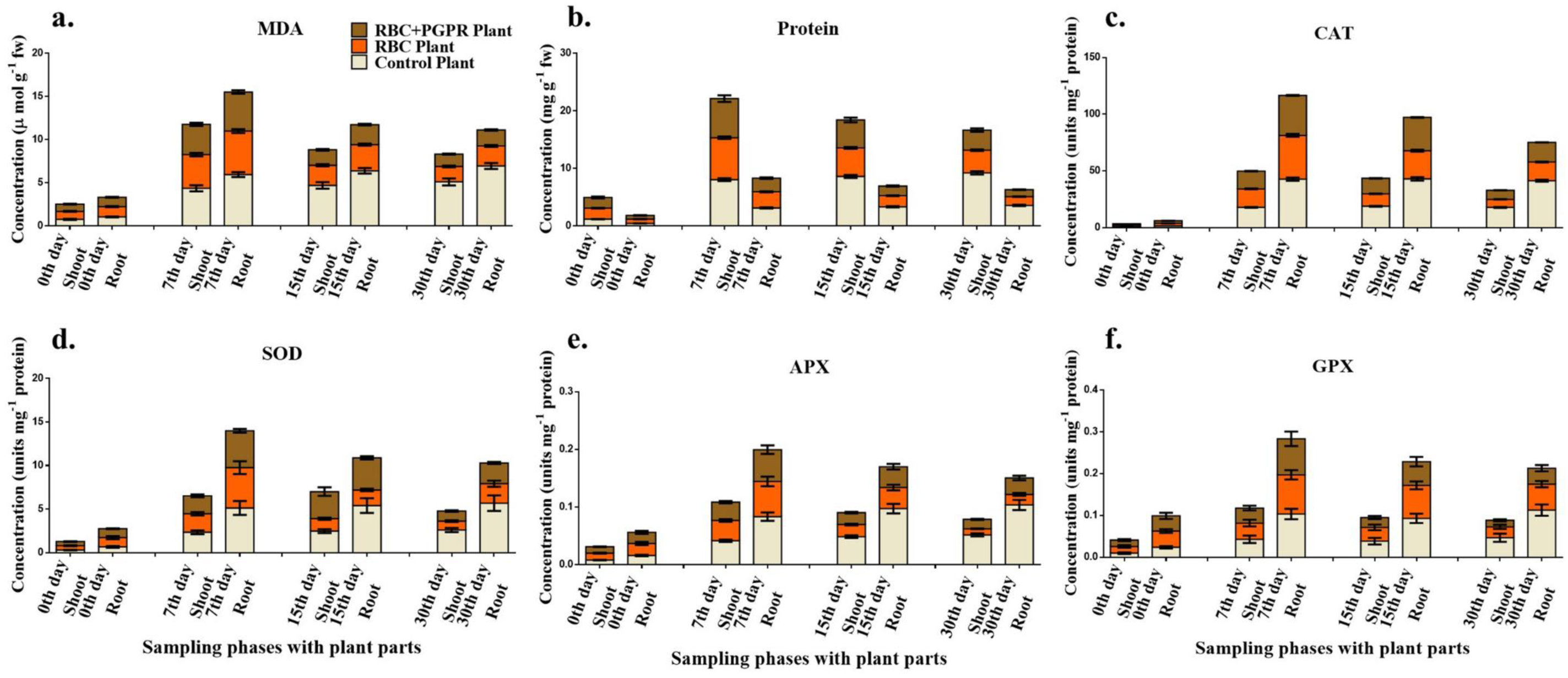
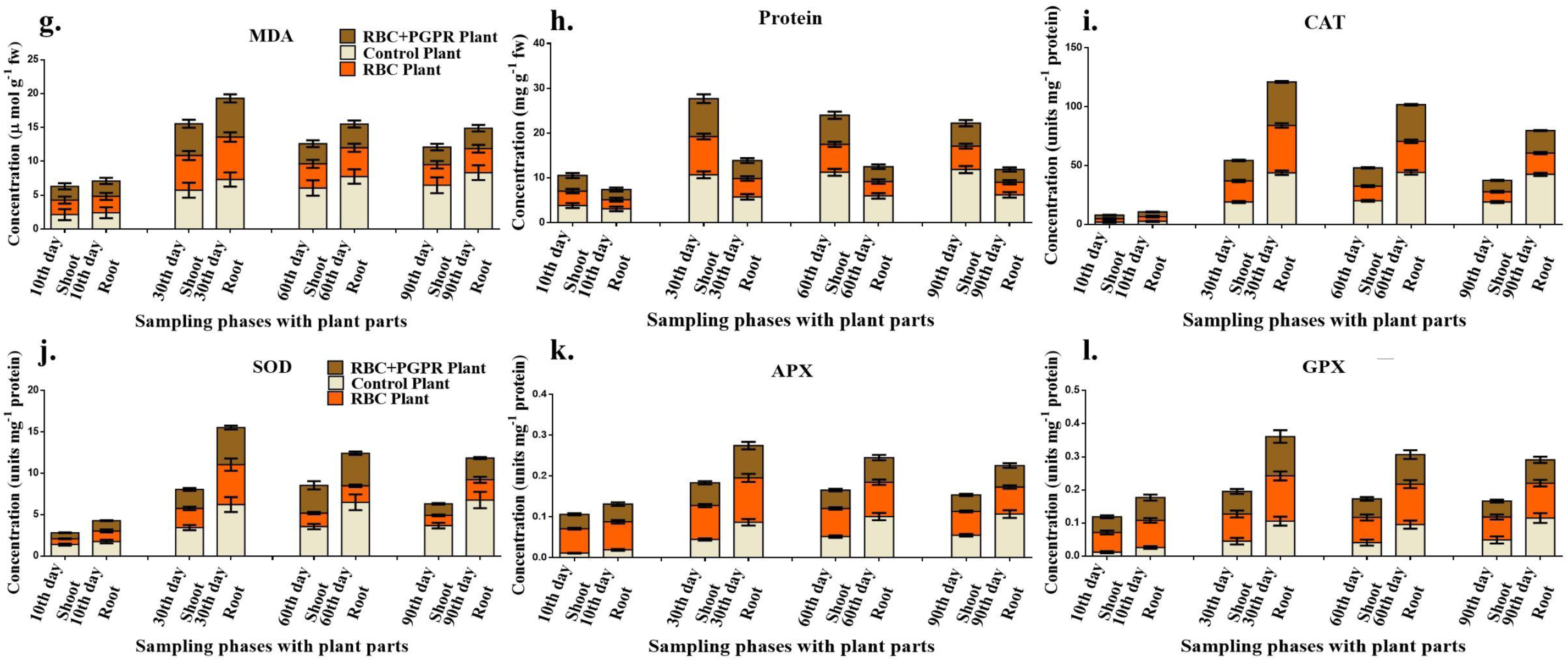
Rice plant antioxidant biochemical analyses towards As stress and after conjugate treatments in pot plants (a-f) and field plants (g-l). Here, 0^th^ day rice seedlings were also considered to compare the enzymatic content with time and treatments. The data are the average of triplicate analysis with shown SD bar followed by the statistical justification by one-way ANOVA at p<0.05.

MDA levels (Figure 2a, g) significantly increased under arsenic stress in control plants, peaking at later growth stages (30th day in pots; 90th day in field), indicating enhanced lipid peroxidation due to ROS accumulation (Finnegan & Chen, 2012). Both RBC and RBC+PGPR treatments markedly reduced MDA content at all time points, with the lowest levels consistently observed in the RBC+PGPR plants, reflecting a synergistic alleviation of oxidative membrane damage. This aligns with prior reports showing that biochar enhances antioxidant defense by adsorbing arsenic and limiting its mobility (Beesley et al., 2011), while PGPR promote As detoxification via arsenate reductase activity and ACC deaminase expression (Rajkumar et al., 2012). Protein content (Figure 2b, h) declined significantly in As-stressed control plants—especially in later stages—suggesting As-induced metabolic inhibition and protein degradation. However, both RBC and RBC+PGPR treatments improved protein levels across all plant parts and time points, with the RBC+PGPR combination yielding the highest recovery. This suggests enhanced nitrogen assimilation and de novo protein synthesis, potentially due to PGPR-mediated N-fixation and hormone signaling (Glick, 2014) and biochar’s ability to improve soil nutrient availability (Lehmann et al., 2011). CAT activity (Figure 2c, i) showed a sharp increase in treated plants, particularly under RBC+PGPR, peaking at the 7th–15th day in pots and 30th–60th day in field conditions. CAT catalyzes the conversion of H□O□ to H□O and O□, a crucial detoxification step under oxidative stress. As reported in earlier studies, PGPR such as Bacillus spp. and Pseudomonas spp. upregulate plant CAT expression via stress priming and hormonal signaling (Han & Lee, 2005). RSB likely contributes by reducing As-induced ROS production at the root–soil interface.

SOD (Figure 2d, j), which dismutates superoxide radicals (O□□) into H□O□, also showed elevated activity under RBC and RBC+PGPR treatments. Maximum SOD activity occurred at 7–15 days in pot plants and 30–60 days in field plants, particularly in roots, suggesting robust early-stage ROS neutralization. Biochar’s porous structure supports microbial colonization, indirectly stimulating SOD through PGPR-root interactions (Xu et al., 2013). Both APX (Figure 2e, k) and GPX (Figure 2f, l) showed significantly higher activity in the RBC+PGPR treatments at nearly all time points, with pronounced peaks at the 7th–15th and 30th–60th day intervals, respectively. APX is a key enzyme in the ascorbate-glutathione cycle that detoxifies H□O□ using ascorbate, while GPX utilizes guaiacol to achieve similar functions. These enzymes are crucial for maintaining intracellular redox homeostasis under prolonged stress (Gill & Tuteja, 2010). PGPRs are known to enhance expression of APX/GPX via induced systemic tolerance (IST), while biochar may modulate the availability of micronutrients (e.g., Fe, Mn, Zn), which serve as enzyme cofactors, thereby enhancing enzymatic function (Yadav et al., 2020). Overall, field-grown plants exhibited a more sustained enzymatic response over 90 days compared to short-term pot experiments, suggesting a cumulative, adaptive plant–microbe–biochar interaction. The synergistic effect of PGPR and RSB was consistently superior to RSB alone, indicating that microbial consortia can significantly enhance redox buffering capacity and metabolic resilience under As stress. These observations confirm earlier hypotheses that PGPR–biochar synergy extends beyond As immobilization into molecular-level modulation of stress signaling pathways, enzyme induction, and membrane stability (Liu et al., 2018).

### Physiological changes in the rice plant under RBC-doped PGPR

The Figure 3 presents comparative scanning electron micrographs depicting the effects of arsenic (As) stress and remediation treatments (rice biochar [RBC], and plant growth-promoting rhizobacteria combined with RBC [PGPR-RBC]) on the stomatal status (St) and vascular anatomy (xylem [Xy] and phloem [Ph]) of rice plants, evaluated under both pot and field conditions. Control Plant (a, g): Stomata appear partially open but with clear cellular distortion and a denser, more compact surrounding surface structure. RBC+ Plant (b, h): Slight improvement in stomatal opening and surface integrity is visible compared to control, indicating partial mitigation of As-induced damage. RBC+PGPR Plant (c, i): Stomatal pores are more distinctly open and surrounded by less disrupted epidermal material, signifying enhanced recovery of stomatal function. Control Plant (d, j): Xylem and phloem tissues show collapse or distortion, with smaller, irregular cell walls and limited lumen area, indicating impaired transport and tissue integrity. RBC+ Plant (e, k): There is increased cell wall definition and lumen area, marking partial restoration of normal anatomical structure within vascular tissues. RBC+PGPR Plant (f, l): The most pronounced recovery is evident here. Xylem and phloem appear larger, more regularly shaped, and structurally robust, indicating substantial restoration of conductive tissue organization.

**Figure 3.**
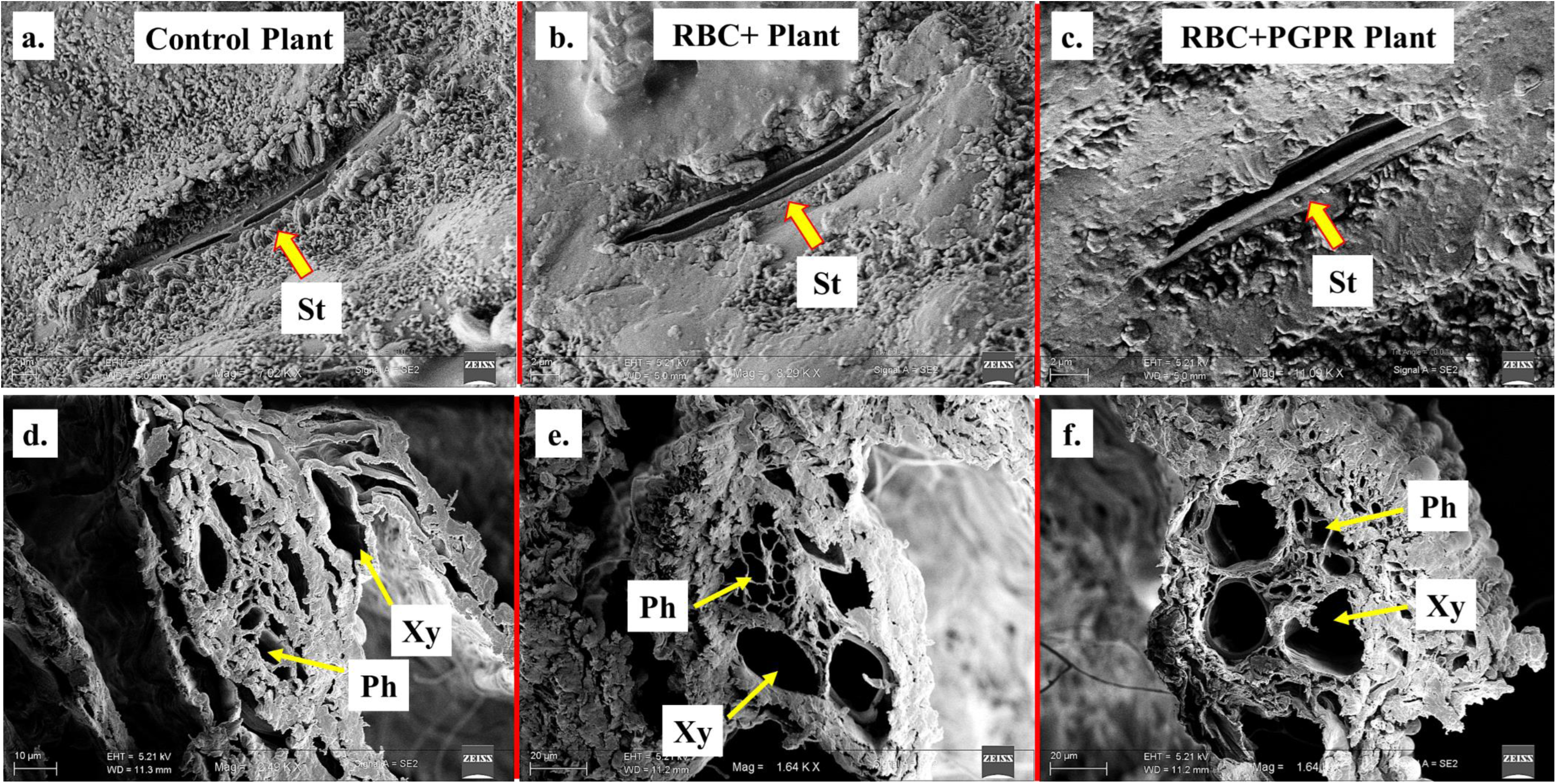
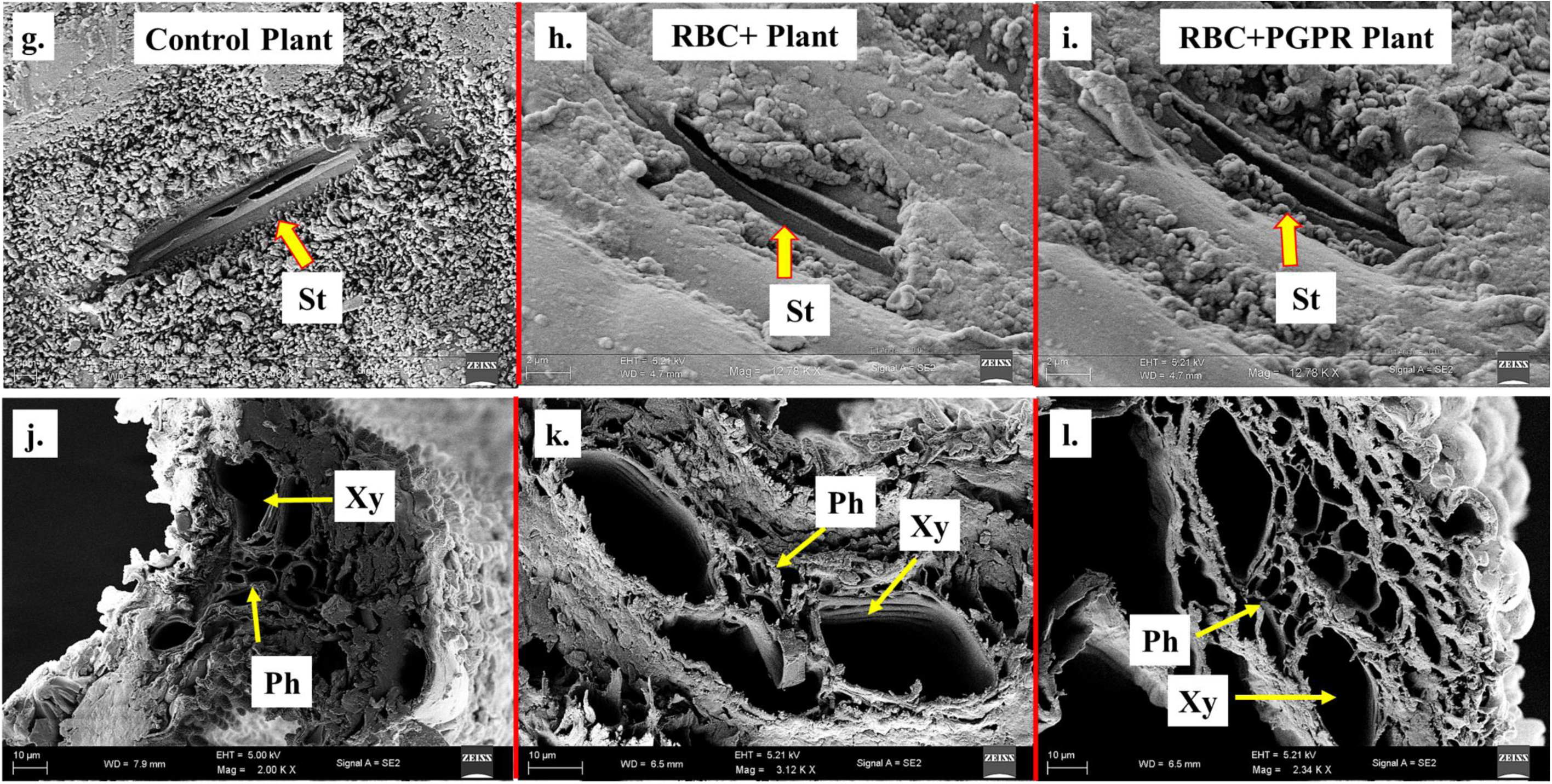
Rice plant stomatal (St) status of gas exchange under As stress and amended setups in both pot-grown (a-c) and field-grown (g-i) plants with vascular anatomy for xylem (Xy) and phloem (Ph) in pot-grown (d-f) and field-grown (j-l) plant shoots.

#### Stomatal Anatomy under As Stress and Remediation

Arsenic (As) stress in rice severely impairs stomatal structure and function, manifesting as deformed guard cells, reduced stomatal apertures, and dense, disorganized epidermal surfaces (see panels a, g for control plants). These observations are consistent with previous reports demonstrating that As toxicity reduces stomatal conductance, inhibits water transport, accumulates osmolytes like proline, and raises membrane permeability—factors that suppress photosynthetic efficiency and plant vitality. The negative correlation between As content and both stomatal conductance and photosynthetic rate in rice has been widely documented (Zhu et al., 2024). Upon introduction of rice biochar, partial restoration of stomatal morphology is evident (panels b, h). Biochar amendments have proven effective in mitigating heavy metal stress by improving soil structure, reducing metal availability, and enhancing physiological processes such as stomatal conductance, transpiration, and water use efficiency. For example, rice husk biochar application has been shown to improve stomatal conductance and gas exchange metrics by stabilizing cellular membranes and reducing oxidative damage (Yuan et al., 2024). The combination of RBC and PGPR (panels c, i) yields the most pronounced improvement: stomatal pores are wider and the epidermal surface appears less damaged, reflecting a high degree of anatomical and functional recovery. This synergistic effect stems from biochar’s As immobilization and PGPR’s ability to enhance nutrient uptake, modulate phytohormones, and trigger plant defense responses. PGPR strains also induce changes in plant gene expression related to cell wall architecture, fostering more resilient guard cells and overall enhanced stomatal function (Grover et al., 2013).

#### Vascular Tissue Recovery

Under As stress, both xylem and phloem tissues display notable damage, observed as collapsed walls and shrunken lumens in control samples (d, j). Such vascular disruption significantly impedes water and nutrient transport, adversely affecting rice growth and yield. Application of RBC (e, k) leads to clearer definition of vascular cells and partial restoration of function. Biochar’s impact on improving water and nutrient availability, as well as reducing heavy metal mobility, directly supports vascular tissue maintenance (Rahman et al., 2019). PGPR, particularly when used alongside biochar, amplifies these benefits (f, l), resulting in the most robust xylem and phloem structures. PGPRs reinforce plant tissues through modulation of cell wall components (e.g., lignin, pectin) and improve overall plant systemic physiology, ensuring efficient water and mineral flow even when exposed to As stress (Kant et al., 2023). Comparative analysis between pot-grown (a–f) and field-grown (g–l) plants validates that the observed morphological and anatomical patterns are consistent across laboratory and agricultural field conditions, demonstrating the robustness of these findings.

### Plant growth promotion and enzymatic boost

Metabacillus indicus (AMB4) displays multiple key plant growth-promoting traits including production of indole-3-acetic acid (IAA), gibberellic acid (GA), siderophores, solubilization of phosphate (P), and ACC deaminase activity, as indicated by elevated metabolite concentrations in PGPR traits and RBC+PGPR setups compared to control (Fig. 4a). These metabolites enhance root and shoot development, stress resilience, and nutrient uptake—IAA and GA stimulate cell division and elongation, siderophores improve iron acquisition, solubilized phosphate makes P biologically available, and ACC deaminase mitigates ethylene-induced stress (Glick et al., 2007, Plant Physiol; Liu et al., 2010, Soil Biol Biochem). The ability of Metabacillus indicus to produce these metabolites is consistent with recent findings showing increased rice yield and improved agronomic performance when treated with Metabacillus indicus, including when chemical fertilizer rates were halved (Fatema et al., 2024; K. Fatema et al., 2024; Saha et al., 2016). Panel b details genes and enzyme activities involved in phosphonate and phosphinate metabolism by AMB4, featuring conversion of 3-phosphoenolpyruvate through multiple metabolic intermediates to pentose phosphates and phosphonates, and highlighting C–P bond cleavage enzymes. These pathways are crucial for using phosphonates as alternative phosphorus sources when inorganic P is limited (McGrath et al., 2013). The ability of Metabacillus indicus to catabolize organophosphonates via C–P lyases and related hydrolases is foundational to its environmental versatility and PGP function, enabling enhancement of soil phosphorus cycling, and is corroborated by bacterial genomic surveys indicating widespread phosphonate metabolism genes in soil bacteria (Kornberg et al., 2012; Lo et al., 2024). AMB4 harbors a suite of genes involved in comprehensive nitrogen metabolism, encompassing nitrate/nitrite reduction (assimilatory and dissimilatory), ammonia production (via nifH and other nitrogen cycle genes), and the regulation of key enzymatic steps (nitrate reductase, nitrite reductase, etc.). These activities drive biological nitrogen fixation and transformation, boosting nitrogen supply to plants. The presence of nifH and other nitrogen metabolism genes in Metabacillus indicus enables significant enhancement of rice seed germination and yield in field trials, even at 50% reduced chemical NPK fertilization, confirming its robust nitrogen-fixing ability and contribution to sustainable agriculture (Jha et al., 2013). Sulfur metabolism genes in AMB4 encode enzymes for the stepwise conversion of sulfate to sulfide and intermediates such as adenylyl sulfate, thiosulfate, and 3-phosphoadenylyl sulfate. These pathways facilitate the mineralization and mobilization of sulfur—an essential plant nutrient impacting protein synthesis and redox balance (Gahan & Schmalenberger, 2014). The ability of Metabacillus indicus to catalyze dissimilatory sulfate reduction, sulfur oxidation, and related reactions enhances sulfur bioavailability to plants, supporting healthy growth under diverse soil conditions (Guo et al., 2016; Camacho et al., 2020; Al-Tammar et al., 2022). Metabacillus indicus (AMB4) emerges as a multi-talented plant growth-promoting bacterium, showing coordinated action in key metabolic traits: phytohormone production, nutrient solubilization, nitrogen fixation, phosphonate utilization, and sulfur cycling. The integration of these mechanisms is pivotal for crop resilience, yield stability, and fertilizer use efficiency. Application of AMB4 in rice systems yields robust benefits, including increased grain yield and reduced dependence on chemical NPK fertilizers (Fatema et al., 2024; Al-Tammar et al., 2022). Genomic analyses further underline the prevalence and ecological importance of genes enabling phosphorus, nitrogen, and sulfur metabolism in Metabacillus and related soil bacteria (Kornberg et al., 2012; Lo et al., 2024; Guo et al., 2016). Collectively, these results support the inclusion of Metabacillus indicus as an efficient biofertilizer in sustainable agriculture.

**Figure 4.**
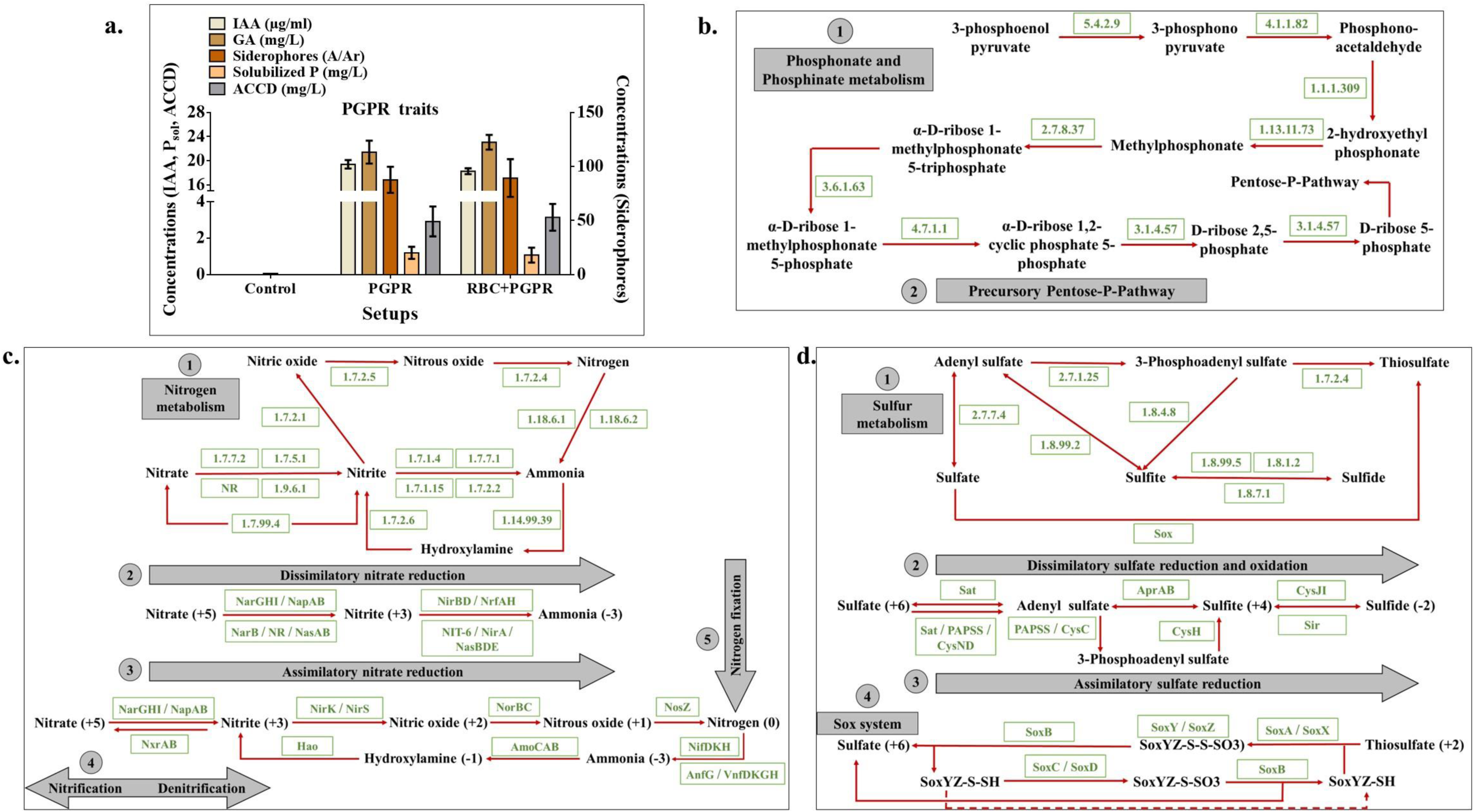
Plant growth-promoting traits in AMB4 (*Metabacillus indicus*), showing plant-required metabolites (a) along with phosphonate metabolism (b), nitrogen metabolism (c) and sulfur metabolism (d).

### Fold change of genes and functional traits with linked networking

Fold-change analysis in Figure 5 revealed significant transcriptional modulation of genes associated with arsenic (As), nitrogen (N), phosphorus (P), and sulfur (S) metabolism, with the blue–red gradient indicating log□ fold-change in activity and lollipop sizes representing gene abundance. Arsenic Metabolism-Exposure to arsenic triggered robust upregulation of arsenic detoxification and transformation genes, including arsR, arsC, arsB, acr3, aox, arsM, and arsH, mediating arsenate reduction, arsenite efflux, methylation, and oxidation (Keren et□al., 2022; Andres et□al., 2016). arsC and arsB exhibited both the largest fold change and highest abundance, suggesting dominant roles in arsenic resistance. These patterns align with environmental genomics reports showing selective pressure for multi-gene ars operons in arsenic-rich soils (Wang et□al., 2024; Stößer et□al., 2023; Xu et□al., 2013). Nitrogen Metabolism-genes encoding high-affinity nitrate/nitrite transporters, assimilatory nitrate reductase (narGHI), nitrite reductases (nirS/K), ammonium transport (amtB), and regulators (glnB/K) were significantly upregulated (Ulrich et□al., 2010; Sobierajska et□al., 2020). The coordinated induction of the GS/GOGAT system indicates efficient ammonium assimilation, consistent with transcriptomic patterns in nitrogen-assimilating PGPR (Chen et al., 2024). Phosphorus Metabolism-Under P-limited conditions, there was strong activation of Pho regulon genes (pstSCAB, phoBR, phoA, phoD), which mediate high-affinity transport and organic phosphate hydrolysis (Santos-Beneit, 2015; Lamarche et□al., 2008). pstS and phoB recorded notable fold changes, consistent with classical phosphorus starvation responses observed in Bacillus and Pseudomonas spp. (Villarreal-Chiu et□al., 2012). Sulfur Metabolism-Sulfur-related genes—soxABCXYZ (sulfur oxidation), dsrAB (dissimilatory sulfite reductase), cys genes (cysteine biosynthesis), and sqr (sulfide:quinone oxidoreductase)—were generally upregulated, especially under S limitation or supplementation (Wolf et□al., 2022; Kies et□al., 2022). Their expression supports enhanced sulfur assimilation and redox balance.

**Figure 5.**
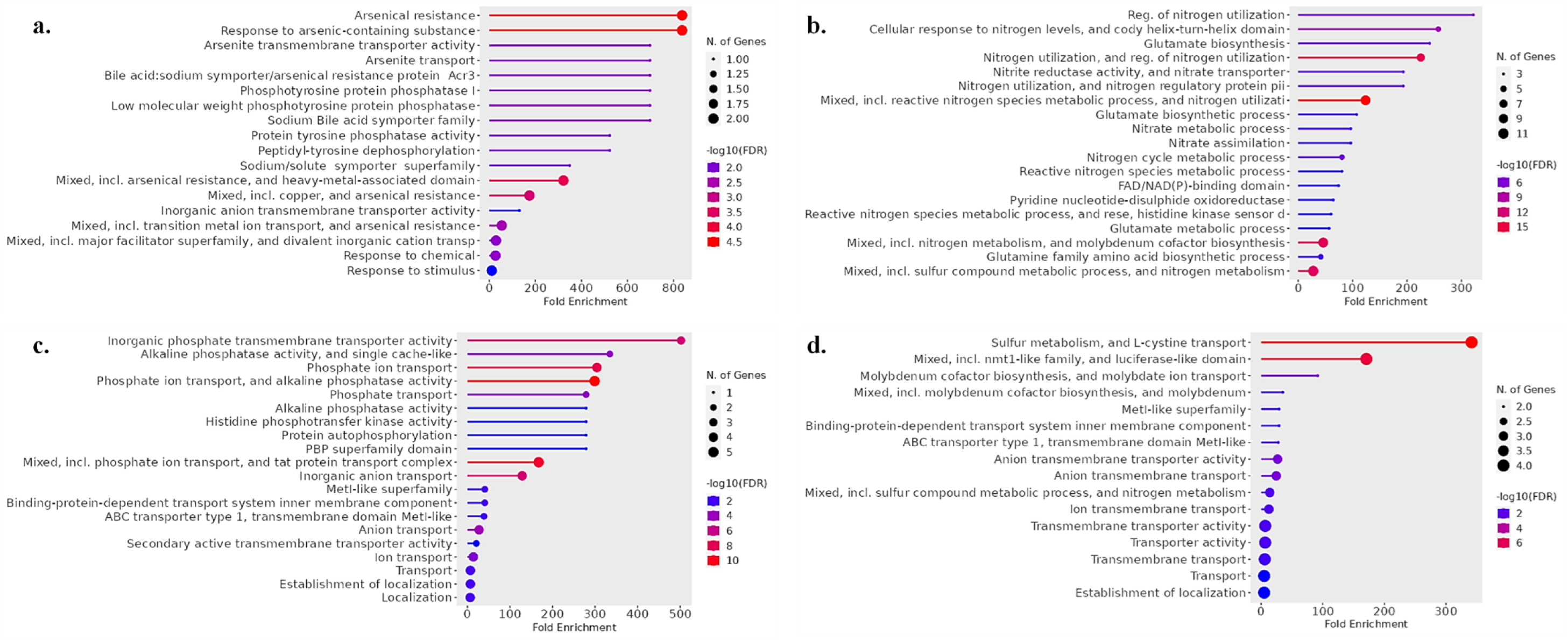
Fold change in the activity of the targeted genes related to their physiological and metabolic expressions. Genes related to As (a), N (b), P (c) and S (d), were analysed for the bacterial metabolic activities and molecular interactions with other related pathways. The fold change of the activity has been shown in the blue-red color code whereas the occurrence of genes has been shown by the size of the circle of this lollipop graph.

Network analysis in Figure 6 mapped the functional linkages among As-, N-, P-, and S-related genes and the proteins they encode, revealing densely connected modules and hub nodes integrating nutrient metabolism with stress adaptation. Arsenic network-Core ars genes linked tightly to oxidative stress regulators such as RpoS, illustrating cross-protection between metal detoxification and general stress responses (Singh et□al., 2025). Phosphorus network-The Pho regulon (phoBR, pstSCAB, phoA/D) formed a central hub, connected to carbon, nitrogen, and sulfur metabolic modules via stress-related pathways (Santos-Beneit, 2015; Lamarche et□al., 2008). Nitrogen network-Nar, nir, glnA, and amtB modules were clustered with glnB/K PII regulators, which bridge nitrogen assimilation, carbon status, and stringent stress response signaling (Ulrich et□al., 2010; Sobierajska et□al., 2020). Sulfur network-sox, dsr, and cys clusters were linked to oxidative stress management modules, highlighting sulfur metabolism’s dual role in nutrient cycling and redox regulation (Wolf et□al., 2011; Kies et□al., 2022). Arsenic detoxification modules shared edges with phosphorus and sulfur pathways, likely reflecting co-regulation under combined metal stress and nutrient limitation (Keren et□al., 2022; Villarreal-Chiu et□al., 2012). The integration through master regulators like RpoS suggests a global transcriptional coordination that optimizes bacterial survival in complex environments. The combined transcriptional and network data from Figures 5 and 6 demonstrate that the bacterium responds to environmental fluctuations through rapid, coordinated activation of As-, P-, N-, and S-metabolic pathways, coupled with global stress regulons. Arsenic genes (ars, aox, arsM) are not isolated modules but part of interconnected networks that engage oxidative stress defense systems (Keren et□al., 2022; Wang et□al., 2024). Phosphorus and nitrogen regulons connect through energy and carbon status sensing via PhoBR and PII proteins, allowing bacteria to adjust metabolism according to nutrient availability (Santos-Beneit, 2015; Ulrich et□al., 2010). Sulfur metabolism serves a dual role—nutrient supply and redox control—bridging energy metabolism with oxidative stress protection (Wolf et□al., 2022; Kies et□al., 2022). These findings align with metagenomic and systems biology studies demonstrating that metabolic flexibility, stress mitigation, and nutrient cycling are intertwined and evolutionarily selected traits in rhizosphere and environmental bacteria (Villarreal-Chiu et□al., 2012; Wolf et□al., 2022). Such integration enhances the bacterium’s ecological fitness and makes it a promising candidate for biofertilizer applications in contaminated or nutrient-poor soils.

**Figure 6.**
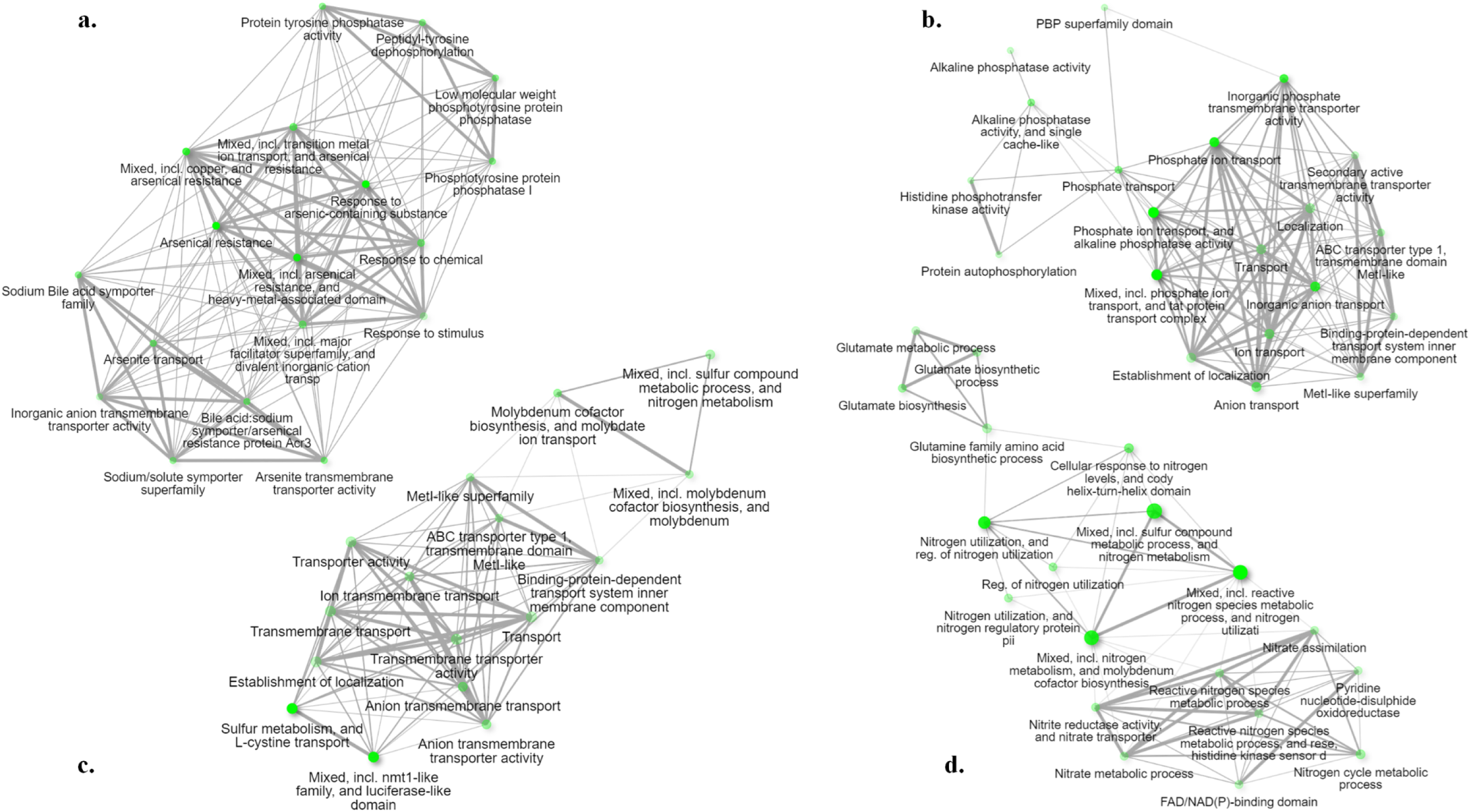
Networking pattern and inter-relationship of As (a), (P (b), N (c) and S (d) related genes and proteins that control major metabolic activities in the bacterial cell along with stress management and nutrient balance.

**Figure 7.**
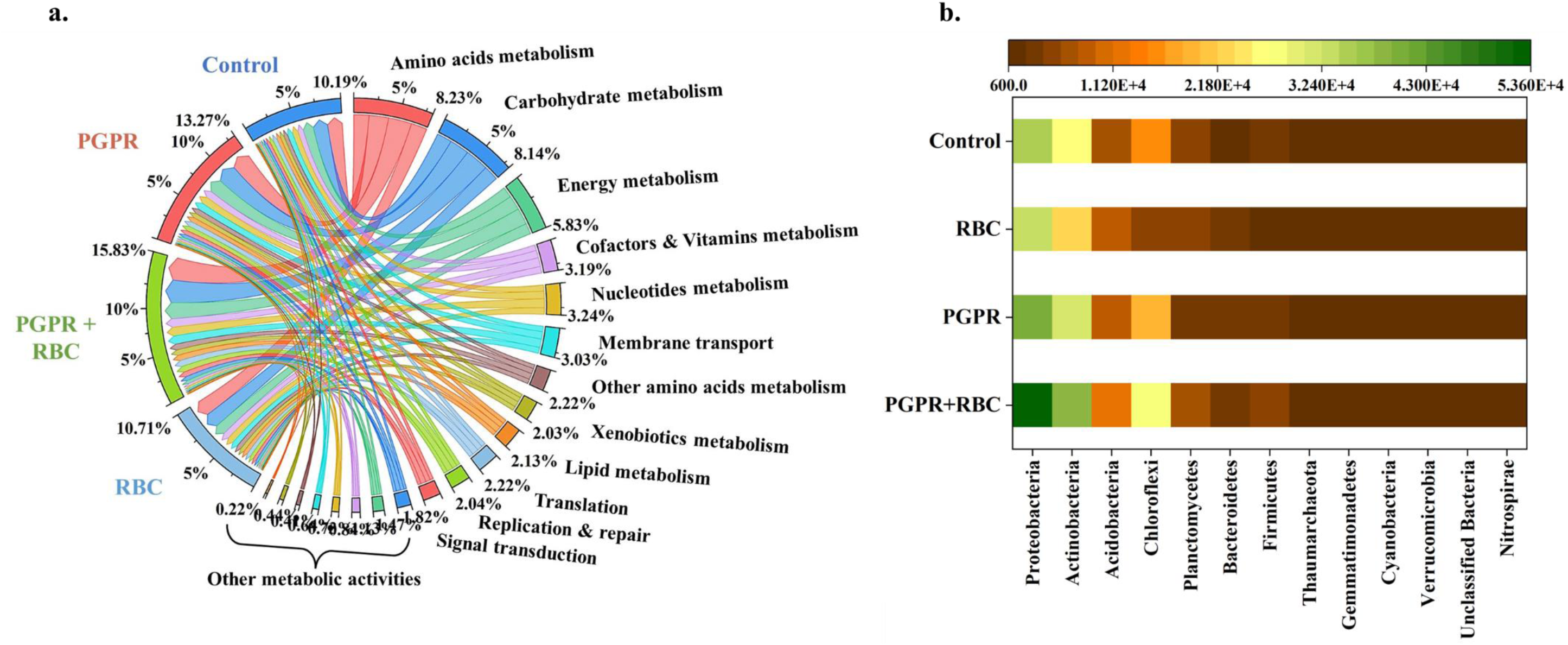
Microbial metabolic activities (a) and phyla distribution (b) under PGPR and RBC application, showing the effect of these amendments on the soil microbial community stabilization and responses.

### Molecular responses of microbiota under differential treatments

This study reveals significant shifts in the metabolic functional potential and taxonomic composition of soil microbial communities in response to amendments with plant growth-promoting rhizobacteria (PGPR) and rice biochar (RBC), individually and in combination.

The functional profiles (Fig. a) demonstrate that the combined PGPR + RBC treatment markedly enhanced the abundance of genes associated with diverse metabolic activities, particularly “other metabolic activities” (15.83%), indicative of elevated microbial metabolic versatility. Singular amendments with either PGPR or RBC also stimulated key pathways compared to the unamended control, underscoring their individual roles in modulating soil microbial functions. The consistently high representation of carbohydrate and amino acid metabolism across all treatments aligns with the crucial functions of soil microbiota in nutrient cycling and organic matter decomposition (Kumar et al., 2020). Importantly, the observed enrichment in xenobiotics metabolism and membrane transport pathways suggests microbial adaptation to altered soil chemistry and potential detoxification processes, which is particularly relevant in contexts of arsenic contamination mitigation (Luo et al., 2018).

Taxonomic analyses (Fig. b) reveal a pronounced increase of Proteobacteria in the PGPR + RBC-treated soils, substantiating the role of this phylum in rhizosphere nutrient transformation and plant growth promotion through mechanisms such as nitrogen fixation and phosphorus solubilization (Garrido-Oter et al., 2018). Additional shifts include enriched Actinobacteria and Acidobacteria populations, taxa known for their involvement in the degradation of complex organic substrates and contribution to soil biogeochemical cycles (Janssen, 2006). These compositional changes signify that the dual amendment effectively restructures the soil microbiome, enhancing both diversity and functional redundancy, which are critical for ecosystem resilience and plant health (Wagg et al., 2019).

Collectively, the integration of PGPR and RBC fosters a microbial community with increased functional capacity and taxonomic composition conducive to nutrient cycling and environmental detoxification. Such synergistic effects likely underpin the improved rice growth and physiological responses observed under these treatments. The amplified metabolic pathways related to amino acid and carbohydrate metabolism further corroborate the role of the microbiome in supporting plant nutrition. Moreover, the engaged xenobiotic degradation systems reflect the community’s adaptability to mitigate soil contaminants, including arsenic, aligning with previous reports on biochar’s role in modulating soil chemistry and microbial detoxification mechanisms (Huang et al., 2017).

This study reinforces the potential of employing PGPR and biochar amendments as integrated strategies to rehabilitate soil microbial function and improve crop productivity, particularly in contaminated or nutrient-poor soils. Future work should leverage metatranscriptomic and metabolomic approaches to dissect the active microbial processes underpinning these community shifts and their direct effects on plant health.

### Microbial classes and their distribution among the treatment setups

Figure□8a presents the total range of microbial class Operational Taxonomic Units (OTUs) detected across the four experimental setups. Variations in OTU richness and distribution reflect how different treatments or environments shape the soil microbial community at the class level (Hartmann et□al., 2015). Generally, management practices such as amendment with plant growth-promoting rhizobacteria (PGPR) or biochar can increase class-level diversity, with the magnitude and pattern of OTU numbers providing insights into overall microbial richness, evenness, and possible dominance among classes (Guo et□al., 2021). Class-level OTU richness is a key metric for assessing ecosystem function, community stability, and response to management or disturbance (Weiss et□al., 2017). In previous studies, higher OTU richness and diversity were directly associated with improved soil health, nutrient cycling, and plant productivity (Li et□al., 2020). Figure□8b shows a scatter matrix plot demonstrating the interdependencies (correlations) between the microbial classes across the setups, with each plot displaying the spread and relationship of OTU counts between pairs of classes. The accompanying correlation coefficients (Pearson’s□r) quantify the strength and direction of these relationships, where positive values indicate co-occurrence, negative values suggest antagonistic or competitive interactions, and near-zero values reflect independence (Durán et□al., 2018; Chen et□al., 2020). High positive correlations (r□>□0.98): Indicative of strongly co-occurring classes, likely sharing similar ecological niches or responding similarly to environmental changes and amendments (Skariah et□al., 2023). Moderate to strong negative correlations (r□∼□–0.56): Reveal competitive or exclusionary relationships among classes, as commonly observed under resource partitioning or niche differentiation (Hartmann et□al., 2015). nter-class variability: Patterns and strengths of these relationships are influenced by soil properties, nutrient availability, amendment type, and external factors such as environmental stressors or plant– microbe interactions (Wolf et□al., 2022). Previous network analyses have shown that positive correlations within bacterial classes tend to dominate, supporting stable community formation, while negative correlations often indicate dynamic competition and potential community restructuring in response to amendments (Chen et□al., 2020; Guo et□al., 2021). The combined range and correlation analysis of microbial class OTUs in Figure□8 reveals not only patterns of diversity and abundance but also the underlying network of interactions that shape community stability and soil ecosystem function (Hartmann et□al., 2015; Li et□al., 2020). Treatments that promote high OTU richness and positive inter-class associations, such as PGPR–biochar co-application, are especially valuable for sustaining soil health and resilience (Guo et□al., 2021; Durán et□al., 2018). Conversely, strong negative correlations may reflect competitive exclusion or stress-induced restructuring, phenomena observed in both natural and managed soils (Chen et□al., 2020; Skariah et□al., 2023). High microbial class diversity with balanced correlations tends to enhance nutrient cycling, pathogen suppression, and ecosystem productivity (Weiss et□al., 2017). Complex correlation structures and network connectivity at the OTU/class level therefore provide a powerful tool for monitoring soil ecosystems, identifying keystone taxa, and predicting responses to agricultural management—paralleling findings from microbial network ecology across terrestrial and aquatic systems (Wolf et□al., 2022).

**Figure 8.**
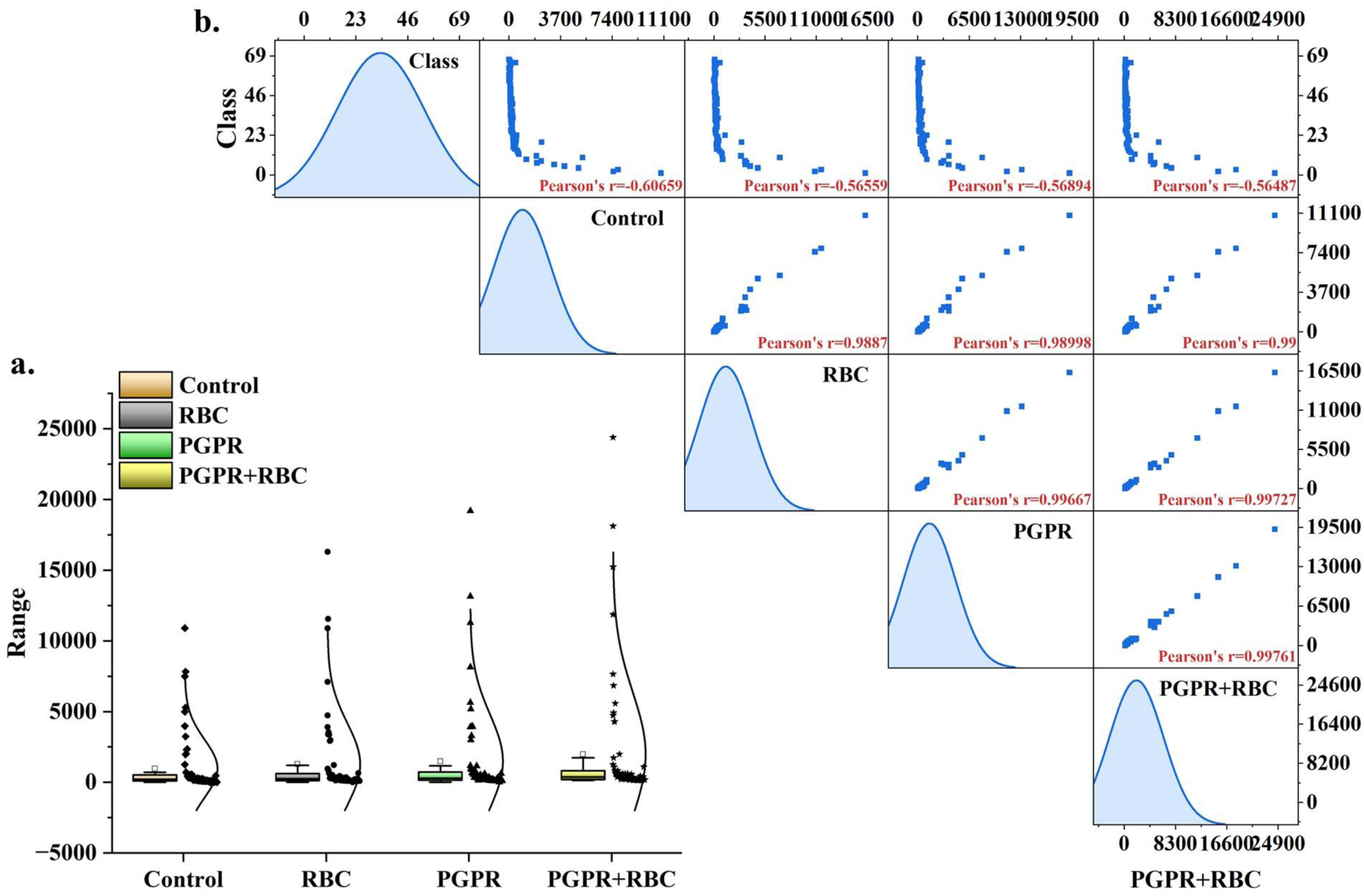
Total range of microbial class OTUs from four setups (a) with their inter-dependent correlation and scattered matrix plot with the correlation coefficient values (b).

## Supplementary Figure Discussion

Figure S1 presents a circular phylogenetic tree constructed from sequence alignments, showing the evolutionary relationships between the selected bacterium AMB4 (Metabacillus indicus) and closely related bacterial species. AMB4 is highlighted on the tree (see the red dot), positioned within a clade containing several Metabacillus and Bacillus species. The clustering demonstrates high sequence similarity and close evolutionary relationship, confirming AMB4’s taxonomic placement within the Metabacillus indicus lineage. Branch support values (often shown as percentages or numbers) provide statistical confidence for each split, with AMB4 forming a robust group with other Metabacillus species. The tree also depicts the broader diversity of related taxa, underlining the genetic distinctiveness of AMB4 compared to both immediate and distant relatives. Such phylogenetic analyses are essential for identifying new strains, confirming taxonomic identity, and inferring the evolutionary history of beneficial bacteria (Fatema et□al., 2024; Gupta et□al., 2020). Figure S2 displays the bioavailability of arsenic (As) in pot soil samples (panel a) and field soil samples (panel b) under different experimental setups (e.g., control, RBC, RBC+PGPR, over time). The y-axis quantifies As bioavailability (mg□kg□¹), while the different colored bars correspond to measurement days (10th, 30th, 60th, 90th day bioavailable As). Control soil: Arsenic bioavailability remains high and relatively stable over time in both pot and field conditions, reflecting limited natural attenuation and continuing risk for plant uptake. RBC (rice biochar) soil: There is a significant reduction in As bioavailability compared to control, which further decreases with time, indicating biochar’s effectiveness in immobilizing arsenic and reducing its plant-available fraction. RBC+PGPR (Plant Growth-Promoting Rhizobacteria) soil: The greatest reduction in As bioavailability is observed with combined biochar and PGPR amendment, often dropping below 2□mg□kg□¹ by later sampling points. This synergy demonstrates enhanced arsenic stabilization and diminished bioavailable pools. Field soils generally show higher initial bioavailable arsenic compared to pot soils, likely due to more complex environmental factors and historical contamination. However, both amendments (biochar, PGPR) are effective in decreasing soil As bioavailability over time in both conditions. Arsenic bioavailability in soil is shaped by soil type, organic matter, pH, iron and aluminum concentrations, and redox status. Amendments such as biochar and PGPR target these mechanisms. Biochar provides strong adsorption sites, decreasing As mobility, while PGPR can further immobilize arsenic through microbial transformation and support for beneficial soil biochemistry. Combination treatments are increasingly recognized for their superior remediation performance in both controlled and field environments (Guo et□al., 2021; Chen et□al., 2020; Hartmann et□al., 2015). This figure directly supports findings from recent work showing that such amendments lower the bioavailable fraction of arsenic in the rhizosphere, reduce the risk from crop uptake, and promote healthier, more resilient soil microbial communities. Figure S3 shows that nutrient concentrations in both pot (a–d) and field (e–h) soils varied significantly across the four sampling phases, with treatments containing rice biochar (RBC) and plant growth-promoting rhizobacteria (PGPR)— particularly in combination—consistently sustaining higher levels of key nutrients such as nitrogen, phosphorus, and potassium compared to unamended controls. This trend was observed in both experimental contexts, indicating that the pot trials effectively modelled field-scale nutrient dynamics. The enhanced nutrient status is attributable to the high sorption capacity and cation exchange potential of biochar, which reduces leaching losses and improves nutrient retention (Lehmann et□al., 2011; Gul et□al., 2015), as well as to PGPR-mediated nutrient solubilization and mobilization via biochemical transformations in the rhizosphere (Ren et□al., 2020; Liu et□al., 2023). The use of one-way ANOVA provided statistical verification of these differences (Virk et□al., 2020), confirming the significance of the observed improvements. Overall, the data support that RBC+PGPR amendments enhance and stabilize soil nutrient content over time, strengthening their application as sustainable soil fertility management strategies in both controlled and field environments. Figure S4 presents arsenic concentrations measured in rice plants grown under different experimental setups in both pots (panel a) and fields (panel b). Across both settings, the data clearly show that rice plants from control (untreated) soils accumulated the highest arsenic concentrations in their tissues. In contrast, plants grown in soils amended with rice biochar (RBC) exhibited significantly lower arsenic uptake, and those from soils with combined RBC and plant growth-promoting rhizobacteria (PGPR) treatments showed the lowest arsenic accumulation overall. This descending trend (control > RBC > RBC+PGPR) is consistent in both pot and field conditions, reflecting the strong ameliorative effect of these soil amendments. The reduction in plant arsenic uptake is attributed to the ability of biochar to adsorb and immobilize arsenic in the soil, thereby reducing its bioavailability, while PGPR further enhance this effect through microbial transformation processes and improved plant health (Hartmann et□al., 2015; Guo et□al., 2021; Ren et□al., 2020). Therefore, Figure S4 supports the conclusion that the combined application of rice biochar and PGPR is an effective strategy to minimize arsenic accumulation in rice plants, promoting safer crop production in contaminated soils. Figure S5 displays the concentrations of key nutrients in rice plant tissues collected from pots (a–d) and field (e–h) across three sampling phases, with results expressed as means□±□SD (triplicate samples) and statistically validated using one-way ANOVA. In both pot and field experiments, plants grown with rice biochar (RBC) and plant growth-promoting rhizobacteria (PGPR)—especially when applied together—consistently exhibited significantly higher tissue levels of essential nutrients such as nitrogen (N), phosphorus (P), potassium (K), and other measured elements throughout all sampling phases compared to control plants. This pattern indicates enhanced nutrient uptake and accumulation in amended treatments, attributable to improved nutrient availability and mobilization provided by biochar’s sorption properties and PGPR’s nutrient-solubilizing and root-stimulating activities (Lehmann et□al., 2011; Liu et□al., 2023; Ren et□al., 2020). The concordance of pot and field results demonstrates the reproducibility and robustness of these effects under varied conditions. Statistical analysis by ANOVA confirmed that the observed increases in plant nutrient concentrations were significant, highlighting the effectiveness of RBC and PGPR— particularly in combination—for optimizing plant nutrition in both controlled and real-world settings (Virk et□al., 2020). These findings align with previous studies demonstrating synergistic enhancement of plant nutrient status under integrated biochar–PGPR management. Figure S6 depicts pre-mature rice plants grown in pots (panel a) and in the field (panel b), highlighting and comparing shoot and root lengths among different experimental setups: control, rice biochar (RBC) amendment, and combined plant growth-promoting rhizobacteria plus biochar (RBC+PGPR). In both experimental conditions, rice plants treated with RBC+PGPR show the longest and most robust shoots and roots, followed by RBC-alone, with control plants displaying the shortest and least developed root and shoot systems. This pattern demonstrates the clear growth-promoting effect of both amendments, especially their combination, which enhances early vegetative development by improving soil structure, nutrient availability, root vigor, and stress resilience (Ren et□al., 2020; Liu et□al., 2023). The results are consistent between pot and field conditions, underscoring the practical reliability of these amendments for sustainable rice production. The mechanisms underlying this improvement include the sorption properties and nutrient retention of biochar and the phytohormone production, nutrient solubilization, and rhizosphere support offered by PGPR (Lehmann et□al., 2011; Backer et□al., 2018). These synergistic effects lead to stronger, healthier root systems and taller shoots—traits directly associated with improved crop establishment and subsequent yield. Figure S7 presents photographs and quantitative data for pot-grown mature rice plants cultivated under different treatments: control, rice biochar (RBC), and the combined application of RBC and plant growth-promoting rhizobacteria (PGPR). The figure displays both tiller numbers (Tn) and panicle numbers (Pn) for each setup, with clear visual and numerical differences across treatments. Control plants exhibit the lowest tiller and panicle numbers (Tn□=□9, Pn□=□37), indicating restricted growth and reduced reproductive output under baseline (untreated) conditions. RBC treatment results in notable improvement (Tn□=□12, Pn□=□59), reflecting the positive impact of biochar on rice vegetative growth and panicle formation—likely due to enhanced soil structure, nutrient availability, and water retention. RBC+PGPR treatment shows the highest values (Tn□=□14, Pn□=□64), demonstrating a synergistic effect that maximizes both tillering and reproductive success. This combination provides superior root-zone nutrient cycling, microbial support, and regulation of plant hormones, resulting in more productive plants. The observed pattern (Control < RBC < RBC+PGPR for both tillers and panicles) closely matches published research showing that biochar amendments increase tiller and panicle production due to better nutrient and moisture dynamics, and that the addition of PGPR further boosts yield-related traits by facilitating nutrient uptake, hormone modulation, and improved physiological vigor (Gul et□al., 2015; Ren et□al., 2020; Liu et□al., 2023; Backer et□al., 2018). The enhancement in tiller and panicle numbers is directly linked to higher overall yield potential and plant health, confirming that integrating RBC and PGPR is highly effective for optimizing rice productivity in contaminated or low-fertility soils. Figure S8 depicts harvested shoot bases of field-grown rice under three treatments—control, rice biochar (RBC), and combined RBC with plant growth-promoting rhizobacteria (PGPR)—with marked circles showing tiller numbers and the relative bulk of shoot material. The greatest tiller density and shoot biomass are clearly observed in the RBC+PGPR treatment, followed by RBC alone, while control plots exhibit visibly fewer tillers and reduced basal shoot mass. This pattern is consistent with experimental and meta-analytic studies showing that rice straw or husk biochar applications significantly increase tiller number and biomass by enhancing soil structure, nutrient retention, and root-zone aeration, especially in low-fertility soils (Kamara et□al., 2015; Singh et□al., 2018; Yi et□al., 2023). The addition of PGPR further accelerates these benefits by promoting nutrient acquisition, root vigor, and phytohormone-mediated growth, as evidenced by greater tiller production and shoot mass in PGPR-treated rice (Abd El-Mageed et□al., 2022; Malik et□al., 2022). Studies confirm that the co-application of biochar and PGPR results in synergistic improvements in tillering, shoot biomass, and yield through complementary effects on soil fertility and rhizosphere microbial dynamics (Gul et□al., 2023; Yi et□al., 2023; Anbuganesan et□al., 2024). Thus, Figure S8 demonstrates that integrated amendment with rice biochar and PGPR is highly effective for maximizing vegetative vigor and tillering in field-grown rice, reinforcing their potential for sustainable yield enhancement in challenging soils. Figure S9 presents phylogenetic trees for arsenic transport and detoxification proteins—ArsA (panel a), ArsB (panel b), and ArsC (panel c)—encoded by the ars operon in AMB4 (Metabacillus indicus) and its closely related bacterial strains. In each panel, the AMB4 protein variants are highlighted among their respective homologs from related genera including Bacillus, Peribacillus, Domibacillus, and Cytobacillus, allowing for a comparative assessment of evolutionary relationships. Across all three trees, the AMB4 proteins (noted in red text) typically cluster within well-supported clades alongside Metabacillus species and certain Bacillus/Peribacillus representatives, reflecting high sequence conservation and likely functional similarity in arsenic export and reduction mechanisms. The branching patterns and bootstrap values (numerical values next to nodes) provide statistical confidence to these evolutionary groupings. The presence of these proteins in AMB4, positioned within conserved clusters, underscores the evolutionary preservation of arsenic resistance strategies via the ars operon among soil and rhizosphere Bacillaceae, and corroborates the critical role of these bacteria in arsenic detoxification across diverse soil environments (Mukhopadhyay et□al., 2002; Andres et□al., 2016; Bröer et□al., 2021). The phylogenetic placement also reveals that, while ArsA and ArsB efflux ATPase and transporter components are closely allied among Metabacillus and Bacillus species, ArsC (arsenate reductase) shows broader distribution and some divergence among taxa (as reflected by branch structure), suggesting possible adaptive evolution for handling varying arsenic pressures in different environments. These trees support the genomic and functional evidence that AMB4 possesses a robust arsenal for arsenic efflux and reduction, inherited from its evolutionary lineage and further supporting its suitability for bioremediation and plant growth promotion in arsenic-affected soils (Zhu et□al., 2017; Marcial Gómez et□al., 2022). Figure S10 illustrates the gene ontology (GO) network for AMB4 (Metabacillus indicus), mapping the molecular landscape of its annotated genes across three major categories: biological processes (a), molecular functions (b), and specific variable activities (c–c1). The diagram highlights yellow-boxed terms, which represent enriched or core nodes with direct parent–child relationships (as indicated by black arrows) that connect child activities or processes to higher-order functional categories at the molecular level. In the biological process network (panel a), AMB4 genes cluster into key metabolic categories such as primary metabolic process, nitrogen compound metabolism, phosphorus metabolic process, and specific organic substance transport and processing, mirrored in their connectivity to specialized metabolic pathways and cellular responses. This reflects AMB4’s expansive ability for nutrient cycling, stress adaptation, and active biotransformation in soil environments (Ashburner et□al., 2000; The Gene Ontology Consortium, 2023). The molecular function category (panel b) shows a central node for catalytic activity branching into a suite of enzymatic functions, including oxidoreductase, hydrolase, transferase, lyase, isomerase, and ligase activities. These functions underpin AMB4’s biochemical repertoire for transforming diverse substrate types, contributing both to plant-beneficial activities and environmental detoxification (Gene Ontology Consortium, 2021; Blake et□al., 2015). The variable activities section (c–c1) provides detailed connectivity for genes involved in smaller or specialized pathways, such as nucleobase, macromolecule, and organophosphonate metabolism. These specific modules, linked hierarchically to parent terms, underscore AMB4’s capacity to metabolize a range of organic and inorganic compounds, supporting its role as a multi-trait plant growth promoter and bioremediator (Camacho et□al., 2020; Fatema et□al., 2024). The marked connectivity (arrows from yellow boxes) emphasizes the gene network’s hierarchical architecture, in which broad functions such as “metabolic process” or “catalytic activity” serve as parent categories integrating more specialized child activities. Such modularity and inter-linkage are characteristic of environmentally versatile microorganisms, facilitating rapid adaptation and multifunctional performance by leveraging interconnected gene pathways (Ashburner et□al., 2000; Camacho et□al., 2020). Figure S11 presents the results of machine learning–based modelling applied to soil microbial community data using K-means cluster analysis (panels a–b) and principal component analysis (PCA, panels c–d) across four experimental setups. In panels a and b, K-means clustering assigns samples into three well-defined clusters based on microbial community composition, as indicated by the separation of initial and final cluster centers and the allocation of observations. This method partitions samples by maximizing similarities within clusters and minimizing distances between clusters, thus revealing distinct community states linked to experimental treatments or environmental conditions (Matchado et□al., 2021). The average and maximum distances in the cluster summary tables also reflect the internal variation and stability of each cluster—lower within-cluster distances suggest greater homogeneity among community types, a hallmark of robust clustering performance. In panels c and d, PCA reduces the high-dimensional microbial abundance data to a two-dimensional plot, where the first principal component (PC1), explaining 99.5% of the variance, distinctly separates the four groups. The relatively small contribution of PC2 (0.36%) highlights that most of the community variation is explained by a single environmental or experimental gradient. The overlaid cluster assignments (panel b) and the PCA biplot with sample and taxa vectors (panel d) visually confirm clear separation and grouping of microbial communities according to their response to different amendments, such as control, RBC, PGPR, and RBC+PGPR treatments. The extracted eigenvector coefficients in panel c identify which variables (microbial features) contribute most to the observed variance, informing the biological processes underlying community differentiation (Nishiyama et□al., 2018; Hugerth & Andersson, 2017). The strong accordance between K-means clusters and PCA visualization underscores the reliability and interpretability of the clustering, with machine learning enabling unsupervised identification of underlying patterns, community states, and treatment effects in complex microbiome data. Together, these results demonstrate that K-means clustering and PCA are powerful, complementary approaches for categorizing microbial communities, uncovering treatment-driven patterns, and reducing dimensionality in high-throughput sequencing datasets. Integrating these methods provides a rigorous, interpretable framework for visualizing and classifying the effects of agronomic interventions and environmental gradients on microbial ecology. Figure S12 displays population pyramid plots comparing the distribution of microbial community taxa under three treatments (panels a–c) with control field microbial communities. In these plots, each horizontal bar represents the relative abundance or proportion of specific bacterial and archaeal groups at various taxonomic levels (typically phylum or class), with the left and right sides reflecting their proportion in treatment and control, respectively. The pyramids visualize how different agricultural interventions (e.g., biochar, PGPR, or their combination) shift the overall structure and balance of the soil microbiome. Compared to the control, the treated communities display marked shifts in the abundance of key microbial groups. For example, treatments typically increase the representation of beneficial taxa such as Actinobacteria, Proteobacteria, and Firmicutes while reducing stress-associated or less beneficial groups like Acidobacteria, a pattern consistent with improved soil fertility and ecological status. The pyramids also show that amendments can decrease the dominance of a few groups (i.e., flatten the pyramid), supporting greater evenness and diversity within the microbial community. These structural shifts indicate heightened microbial resilience, functionality, and ecological stability, aligning with meta-analyses showing that biochar and PGPR amendments reconfigure soil microbial communities toward states associated with enhanced nutrient cycling, disease suppression, and higher productivity (Ren et□al., 2020; Yin et□al., 2021; Guan et□al., 2023). Population pyramid plots are increasingly used in microbial ecology to visually convey changes in richness, evenness, and community structure after environmental or management interventions (Vos et□al., 2017; Banerjee et□al., 2018). The observed changes in pyramid profiles here indicate a successful recruitment and stabilization of beneficial, functionally important microorganisms under amendment treatments.

## Acknowledgement

This work was funded by the National Postdoctoral Fellowship scheme, Ministry of Education, Government of India, with a project number PDF/2022/001418/LS and Marie Skłodowska-Curie-UKRI Postdoctoral Fellowship scheme, United Kingdom, with a File number 101152605 and Council reference-EP/Z002664/1. A.M. acknowledge the support from Imperial College London Library during this study and funding for the publication.

## Authors information

### Authors contribution

A.M. conceptualised the project, received funding, performed experiments, and wrote the manuscript; I.K-L., M.B. and T.C. supervised the projects and revised the manuscript.

---

**Figure S1.**
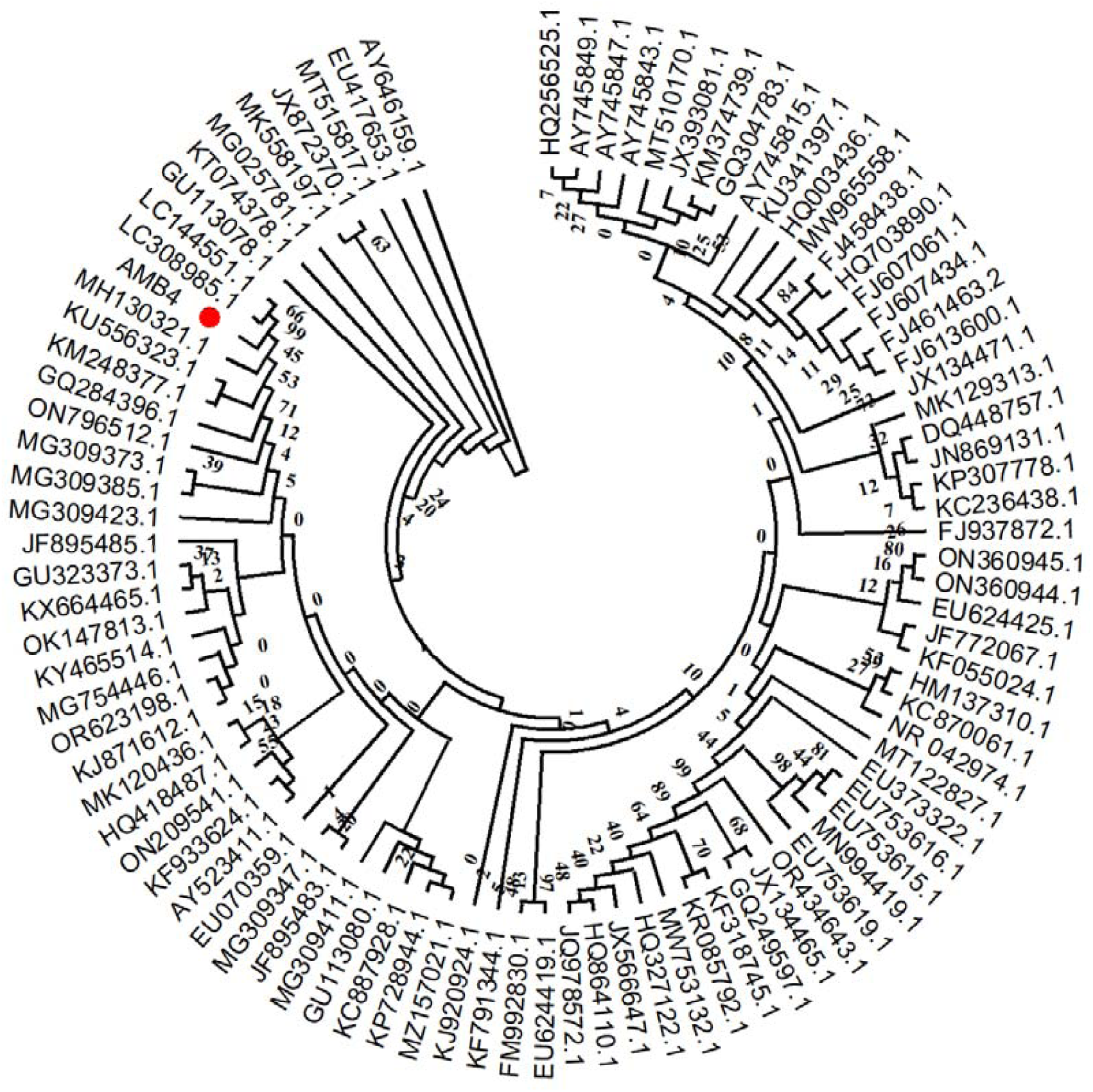
Phylogenic placement of the selected bacterium AMB4 with closely related bacterial species identified from sequence alignments.

**Figure S2.**
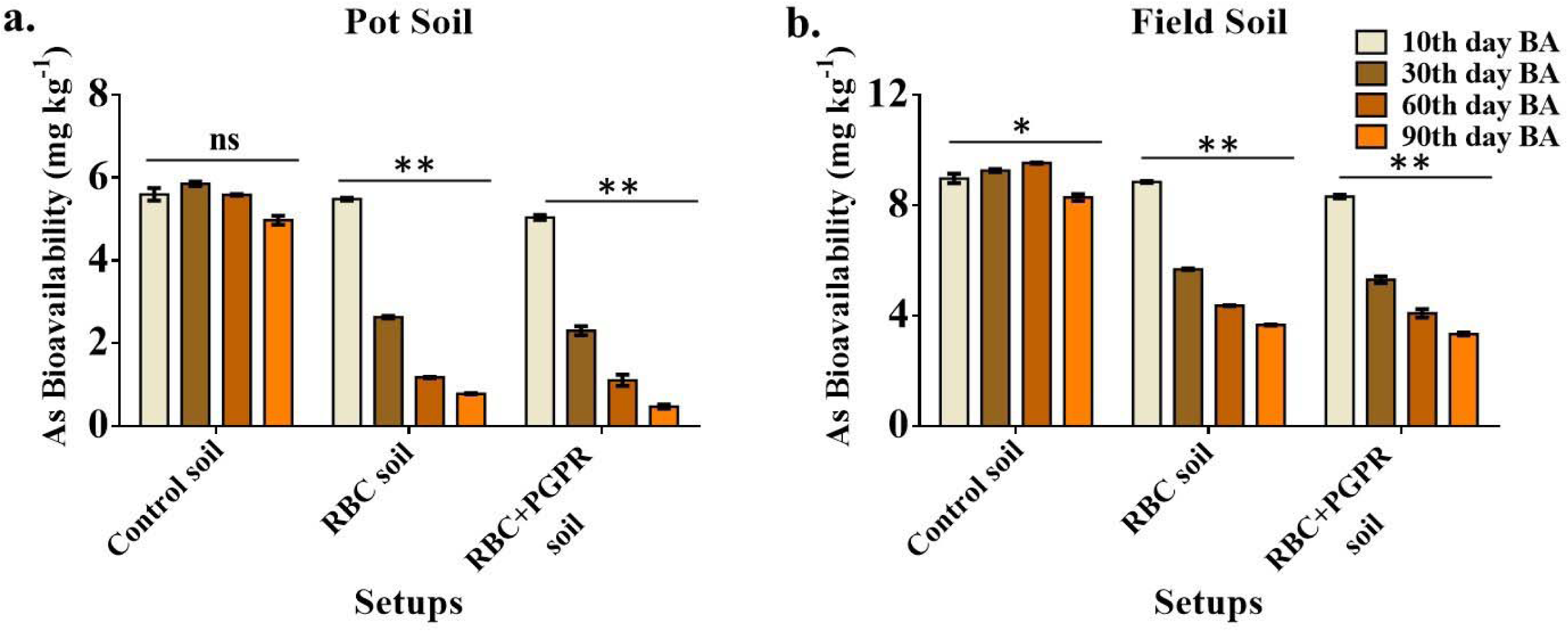
Arsenic bioavailability in pot soil samples (a) and field soil samples (b) with differential setups.

**Figure S3.**
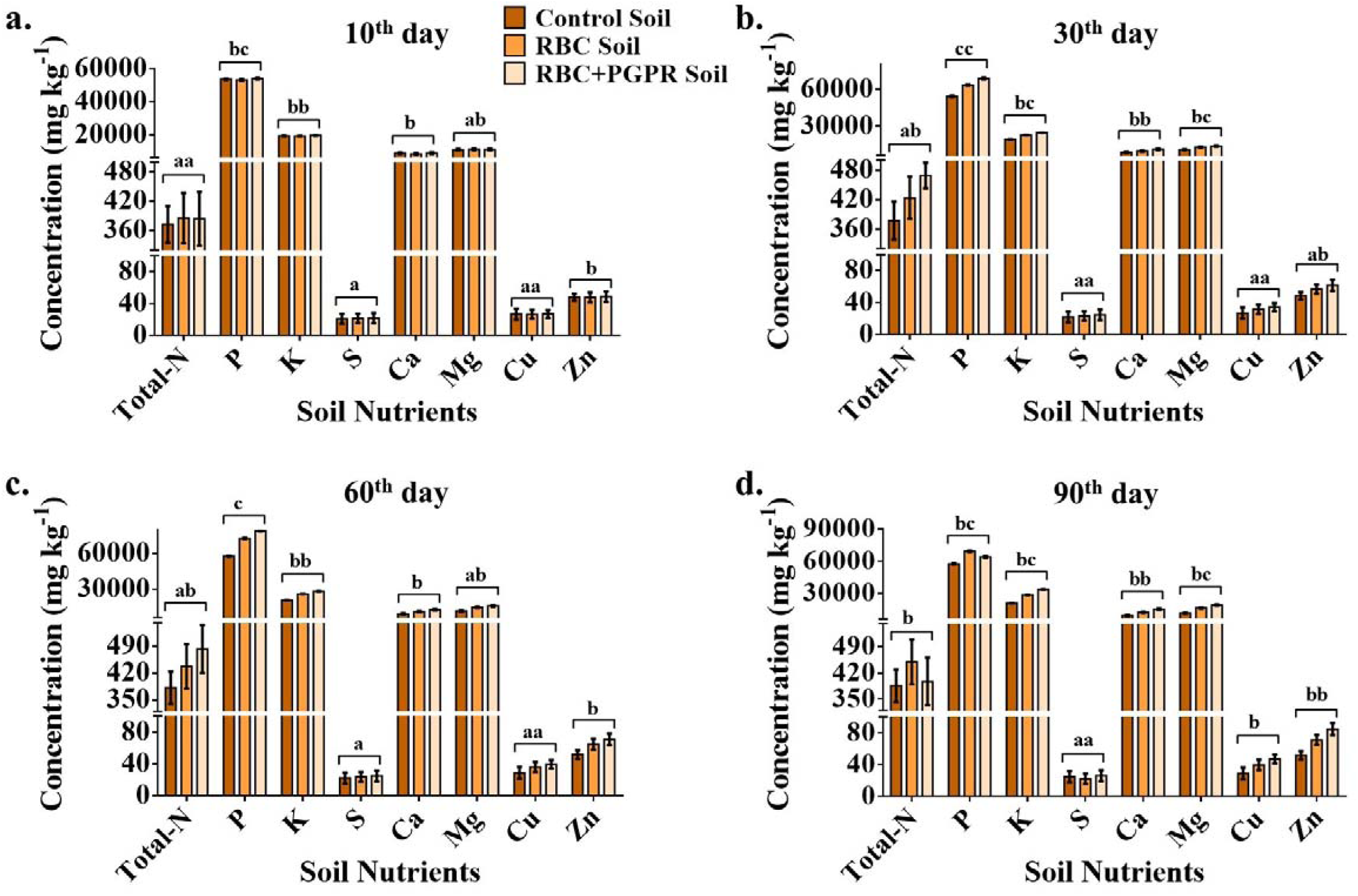
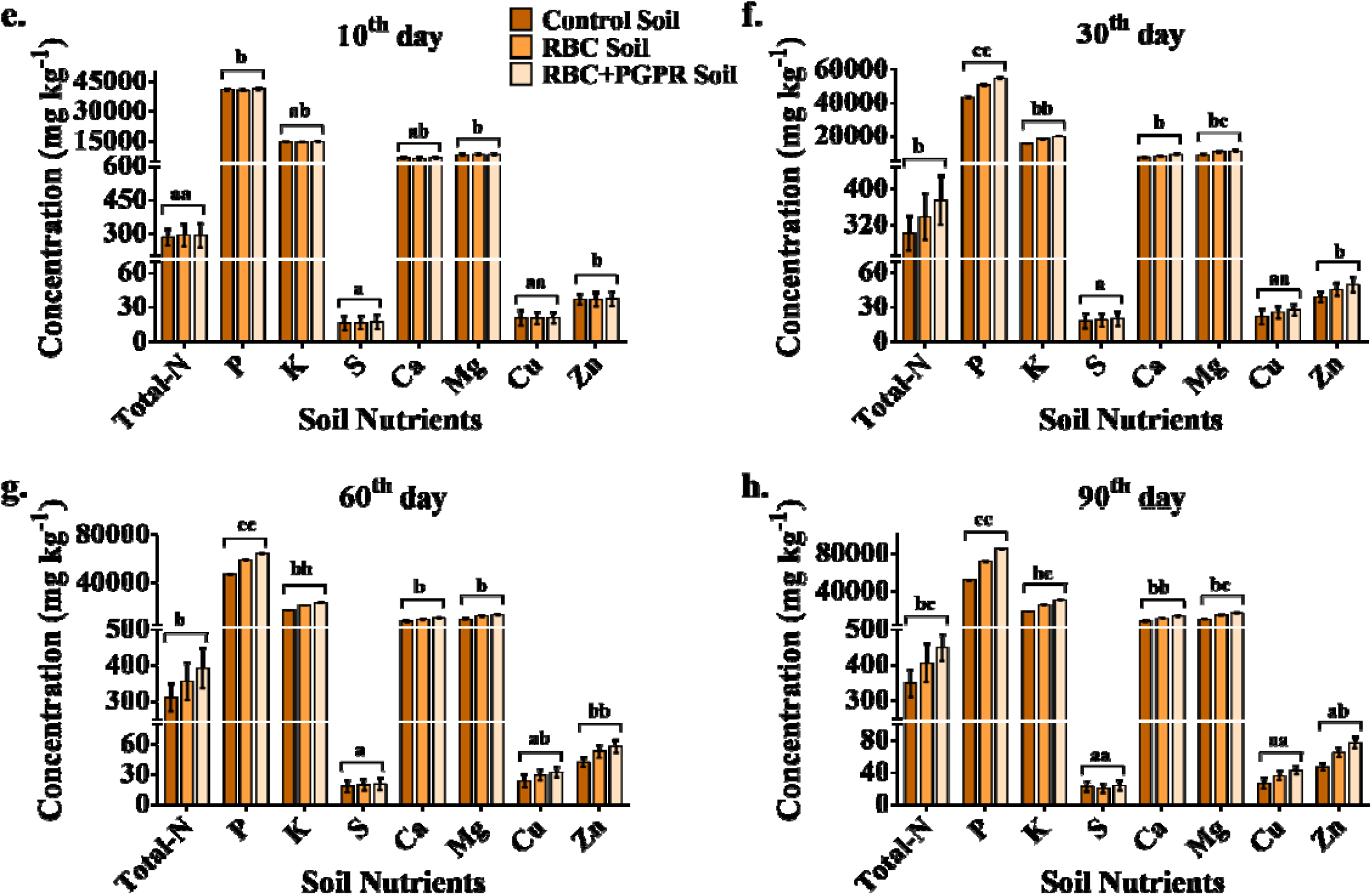
Nutrient content in the pot (a-d) and field (e-h) soil samples at four sampling phases. The data are the average of triplicate analysis with shown SD bar followed by the statistical justification by one-way ANOVA at p<0.05. DMRT was also used for the inter-treatment samples to identify the significant difference, marked by the letter on top.

**Figure S4.**
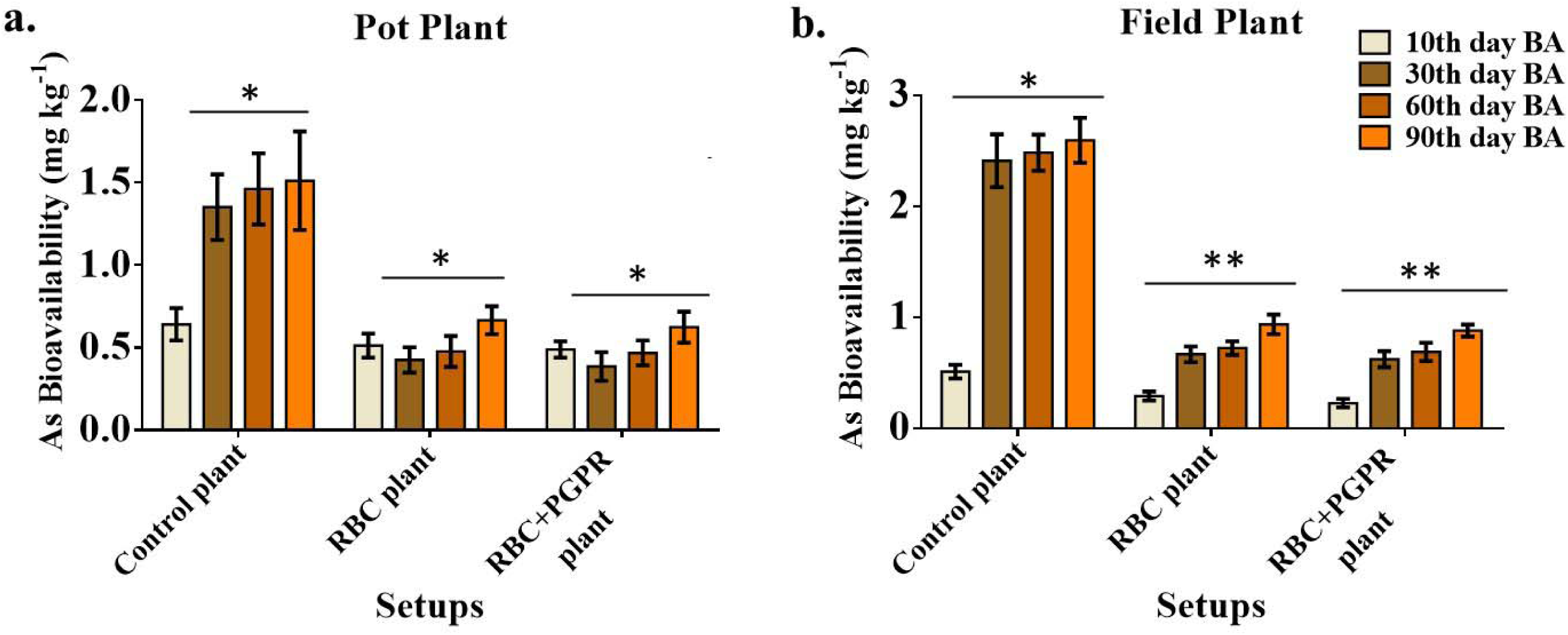
Arsenic concentrations accumulated in rice plants under differential setups in pots (a) and the fields (b).

**Figure S5.**
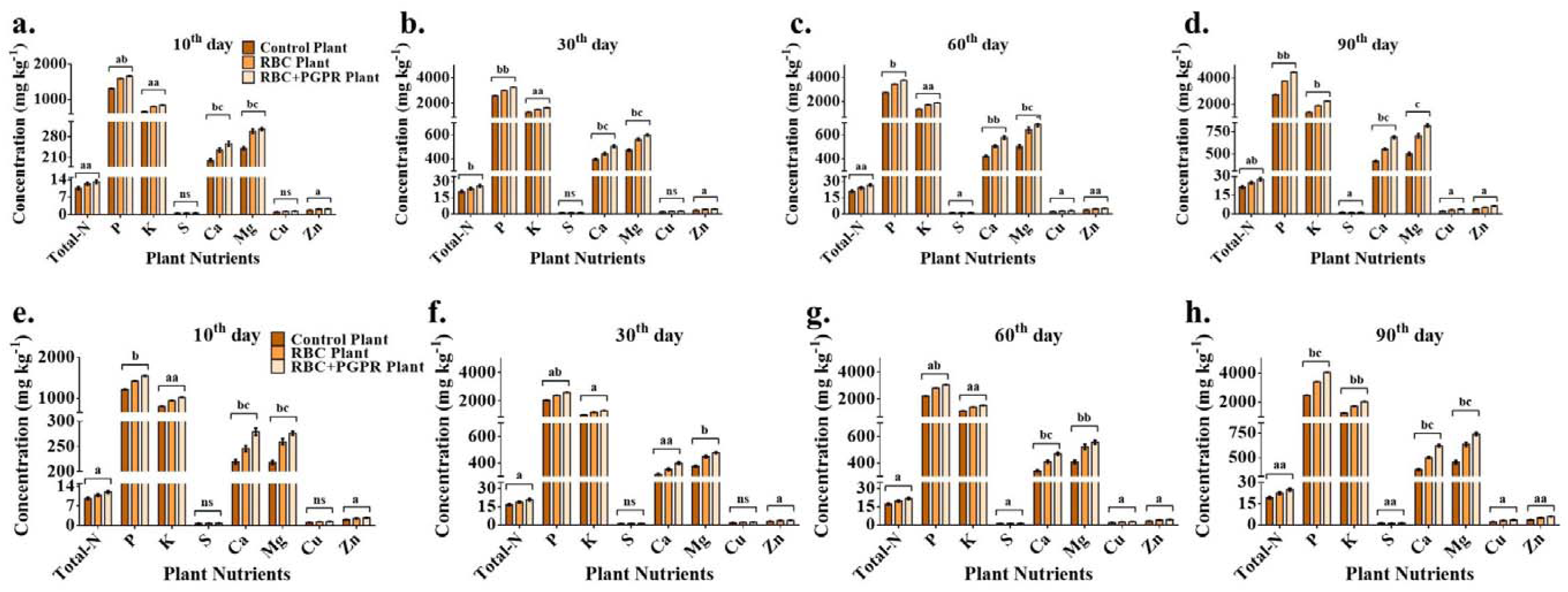
Nutrient content in pot (a-d) and field (e-h) plant samples at three sampling phases. The data are the average of triplicate analysis with shown SD bar followed by the statistical justification by one-way ANOVA at p<0.05. DMRT was also used for the inter-treatment samples to identify the significant difference, marked by the letter on top.

**Figure S6.**
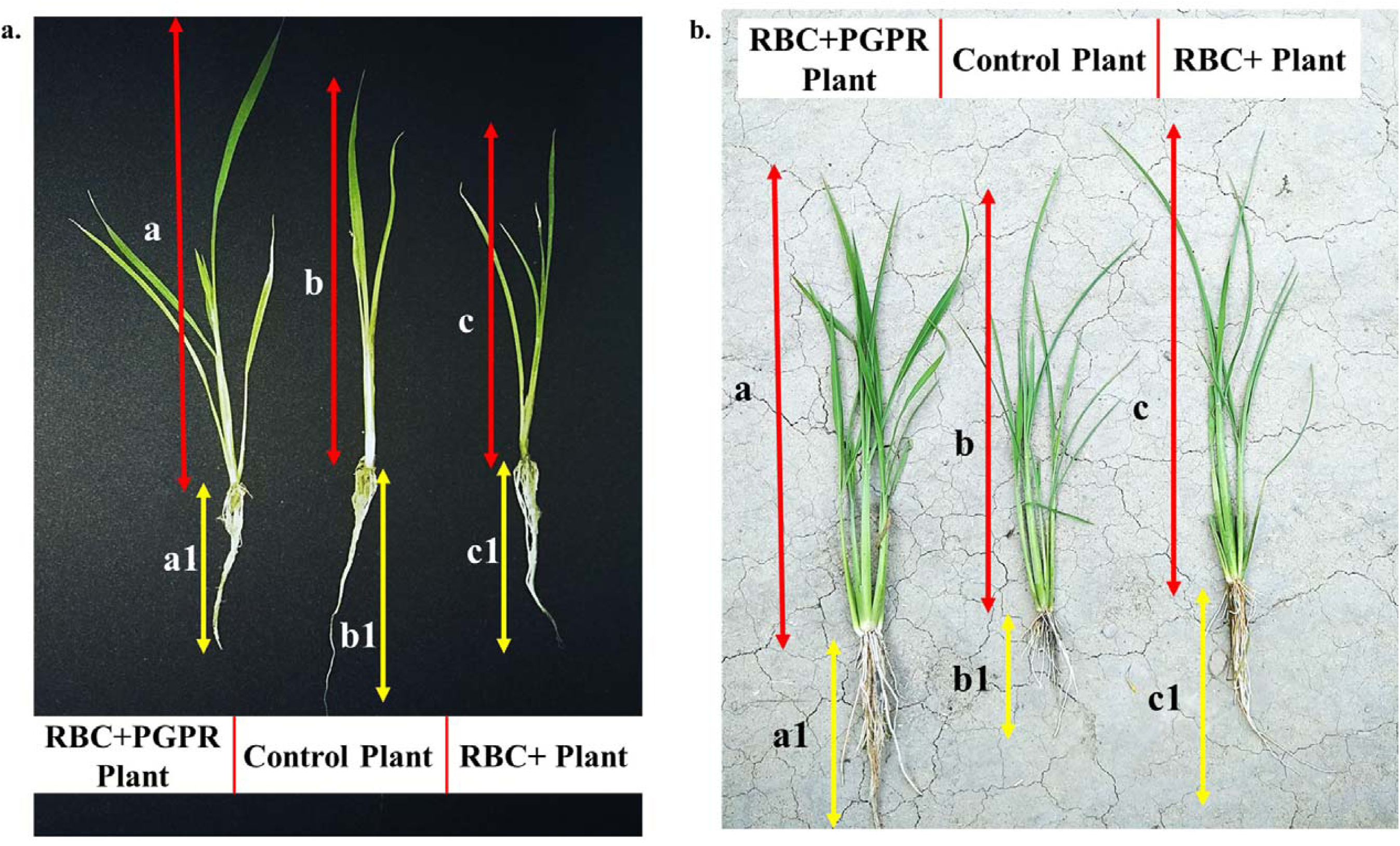
Pot-grown (a) and field-grown (b) pre-mature rice plants with their marked lengths of root and shoot under differential setups.

**Figure S7.**
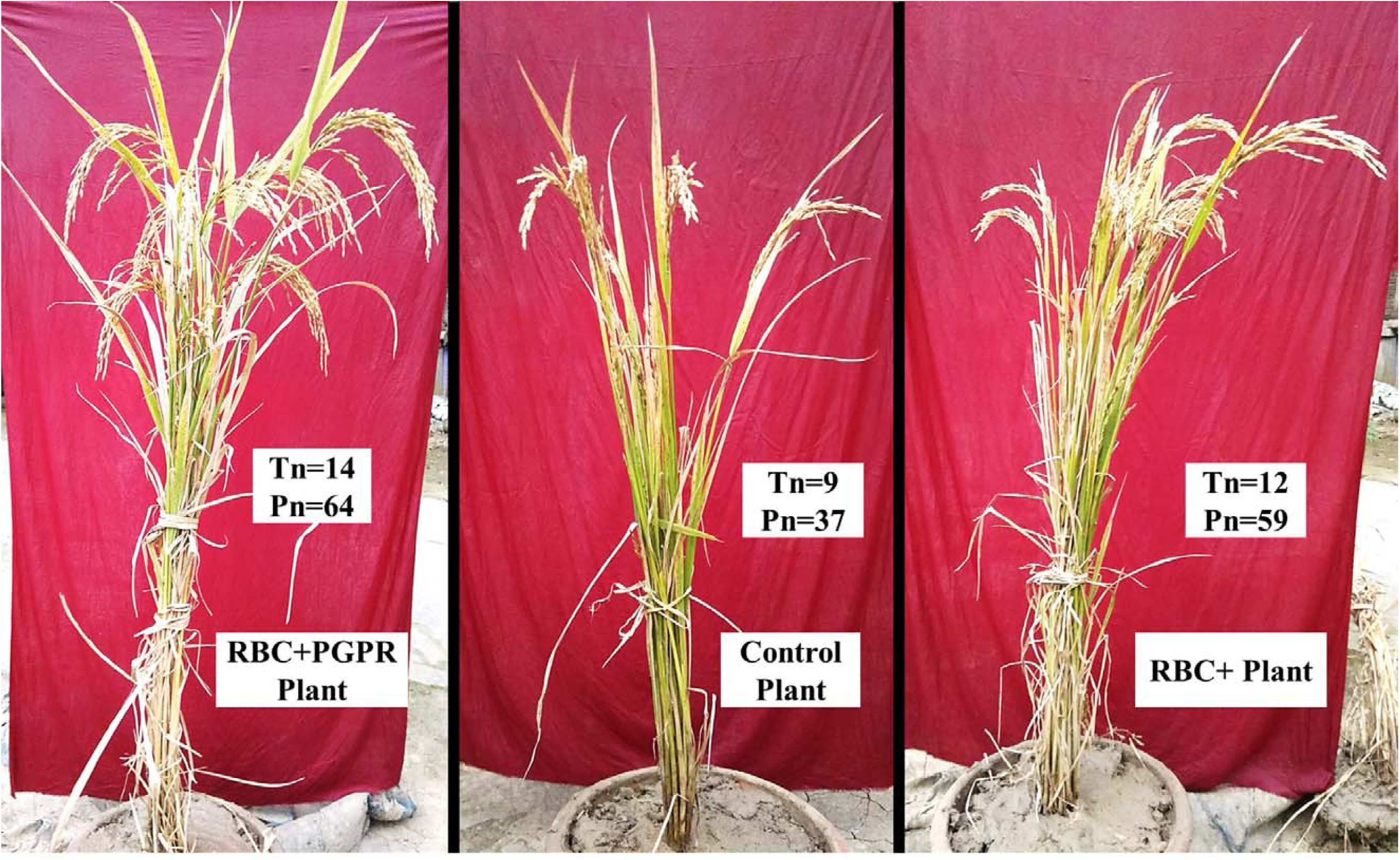
Pot-grown mature rice plants with their tiller numbers (Tn) and panicle numbers (Pn) under differential setups.

**Figure S8.**
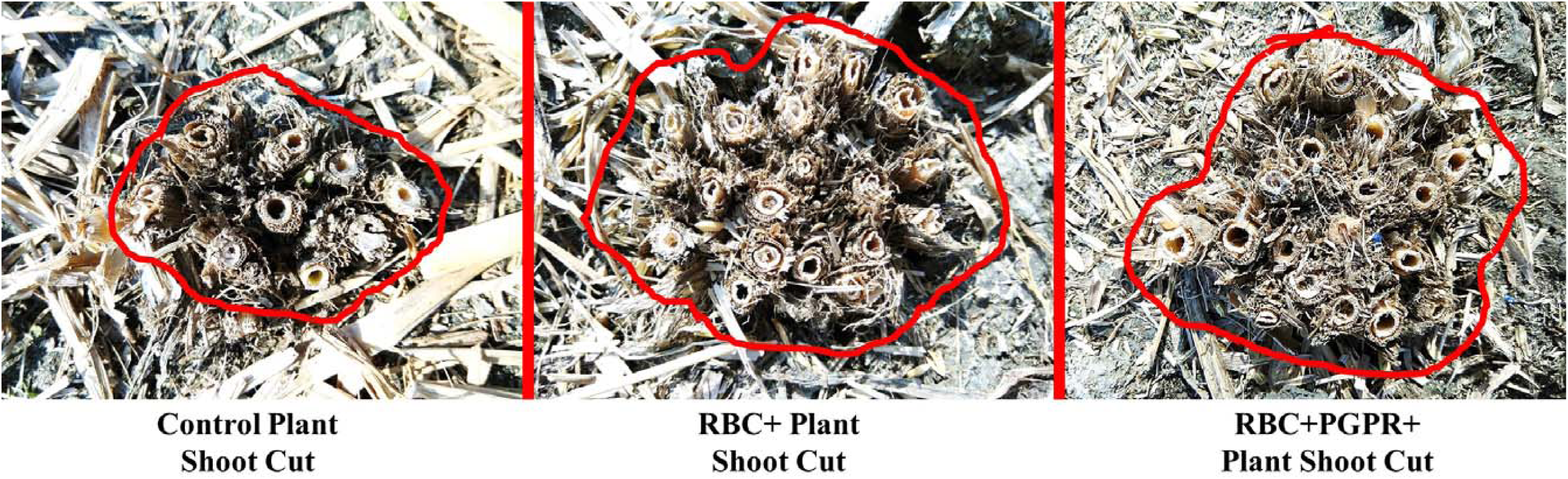
Field-grown rice plant’s cut shoots after harvesting showing bulk growth difference and tiller numbers within marked circles under control and treated setups.

**Figure S9.**
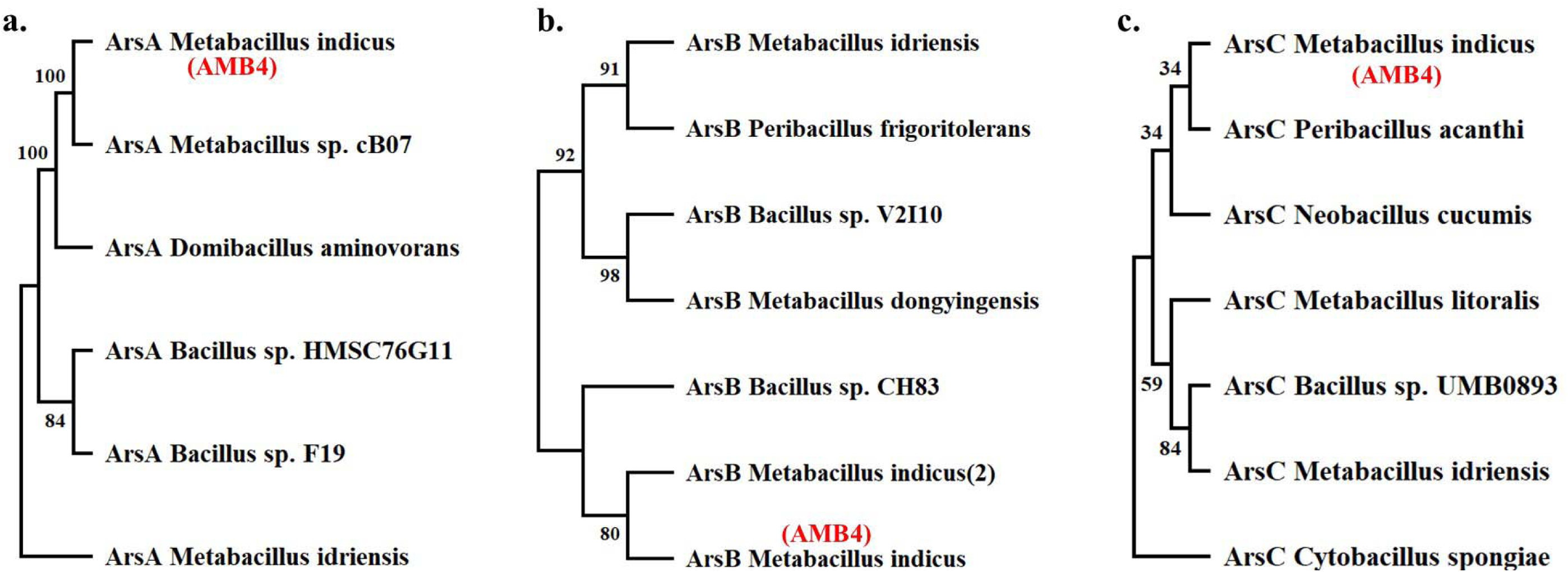
Arsenic transporting gene-responsive proteins ArsA (a), ArsB (b) and ArsC (c) (from *ars* operon) in AMB4 and closely related bacteria with their phylogeny distribution.

**Figure S10.**
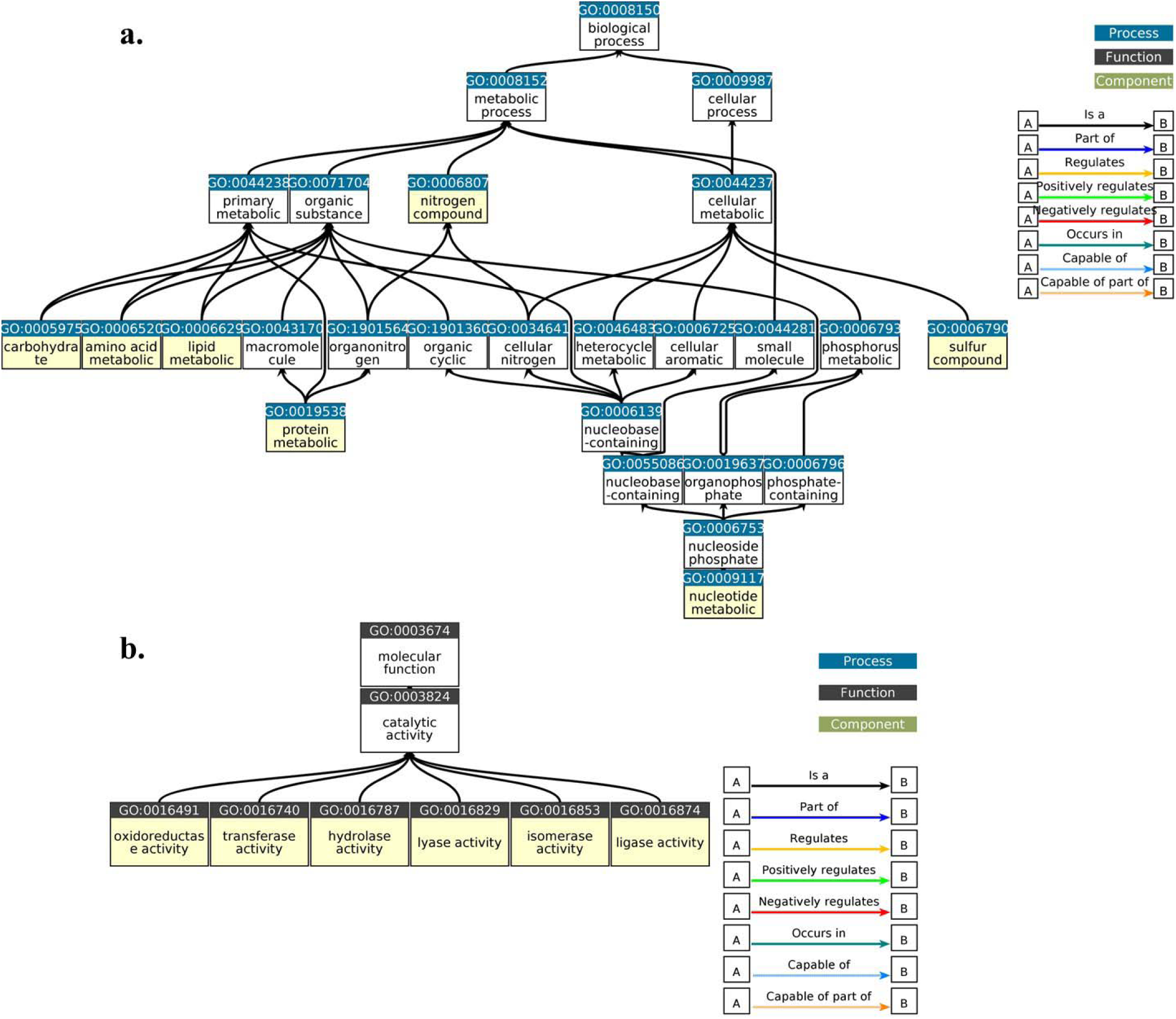
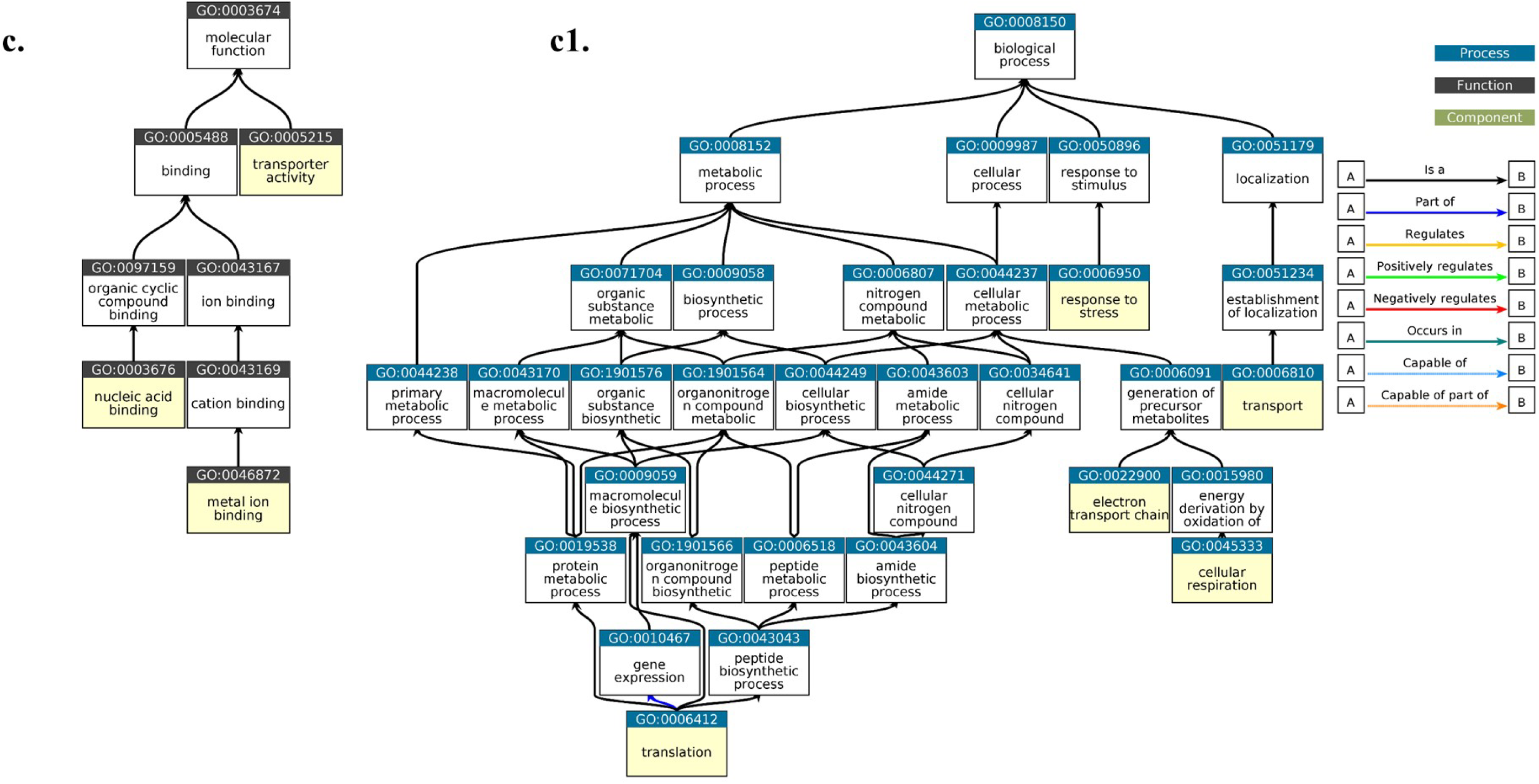
Gene ontological connectivity at the molecular level, showing biological processes (a), molecular functions (b) and variable activities (c-c1) for the AMB4. These inter-connections (specifically within yellow boxes) are marked with the black arrow shown beside indicating their linkage to their parent functions at the molecular level.

**Figure S11.**
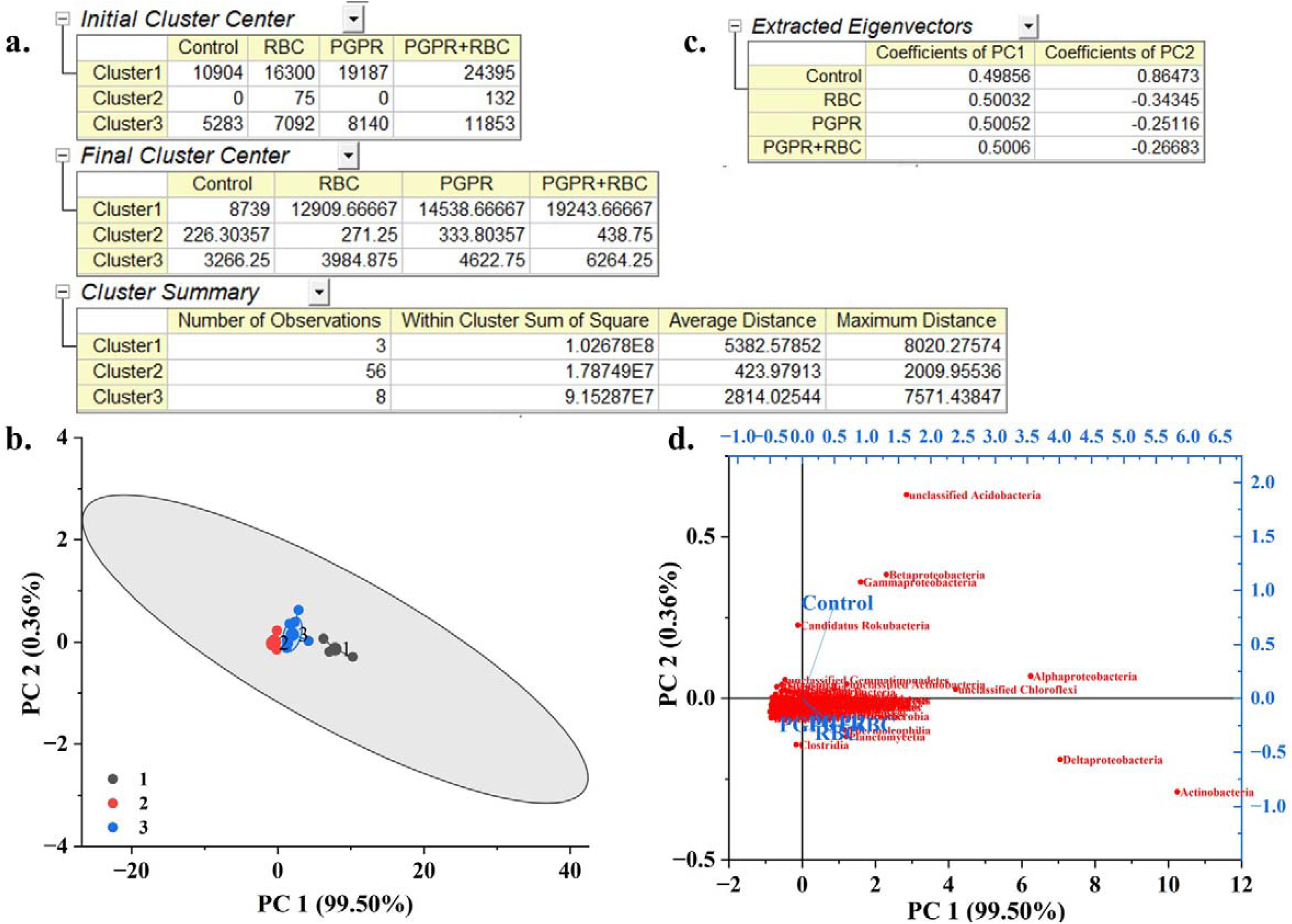
ML-based modelling of K-means cluster analysis (a-b) and PCA (c-d) are performed on microbial community data with four setups.

**Figure S12.**
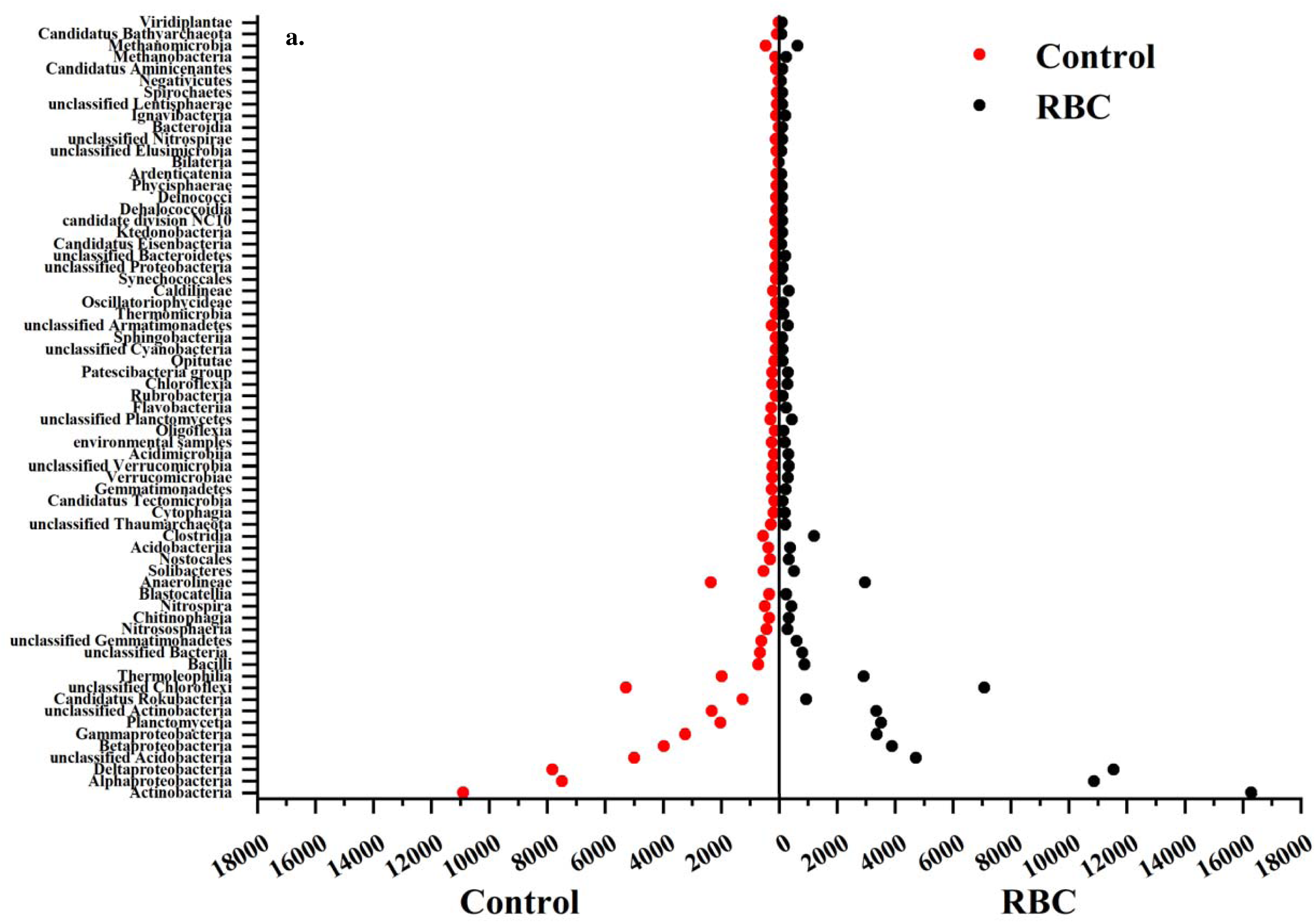
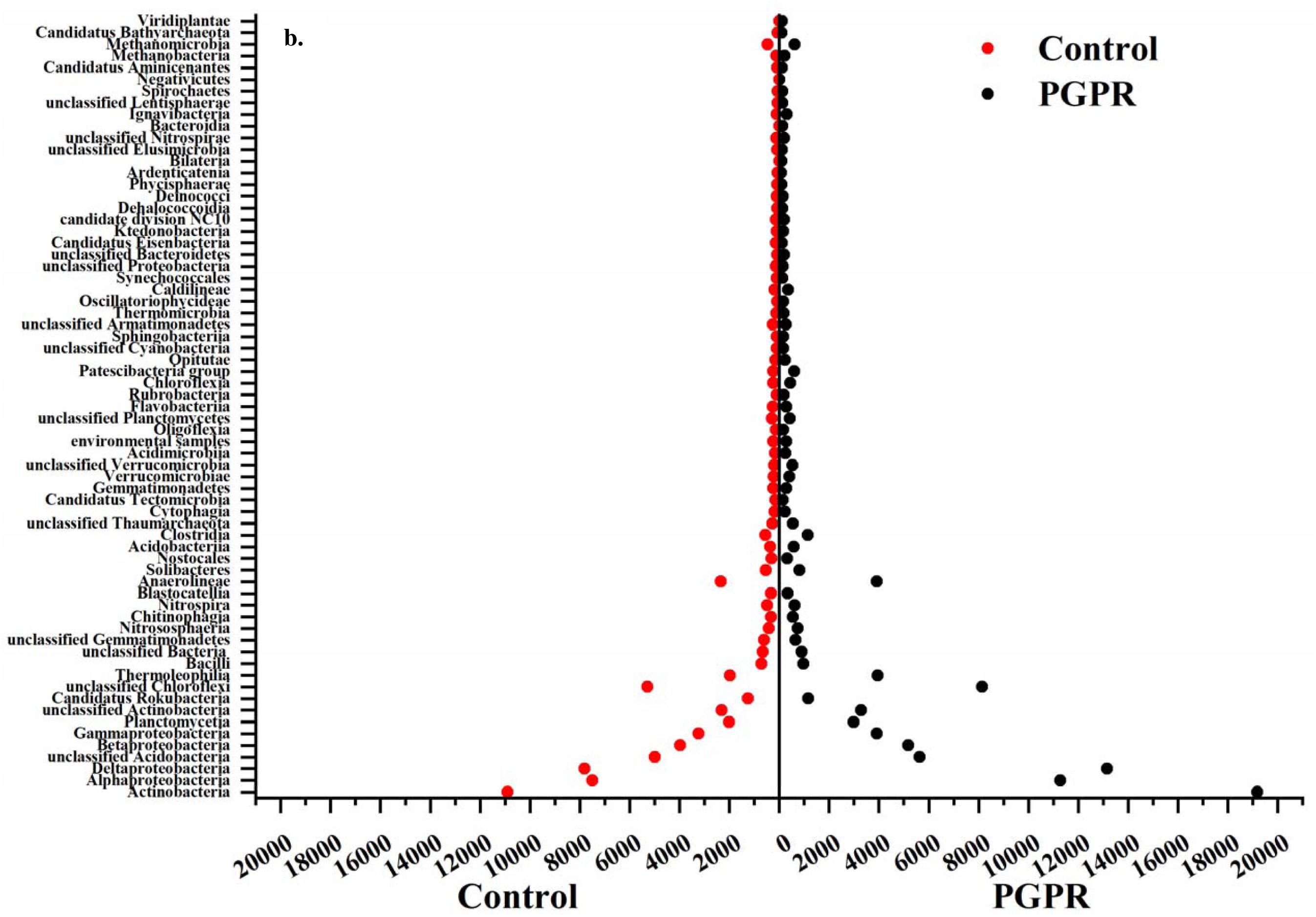
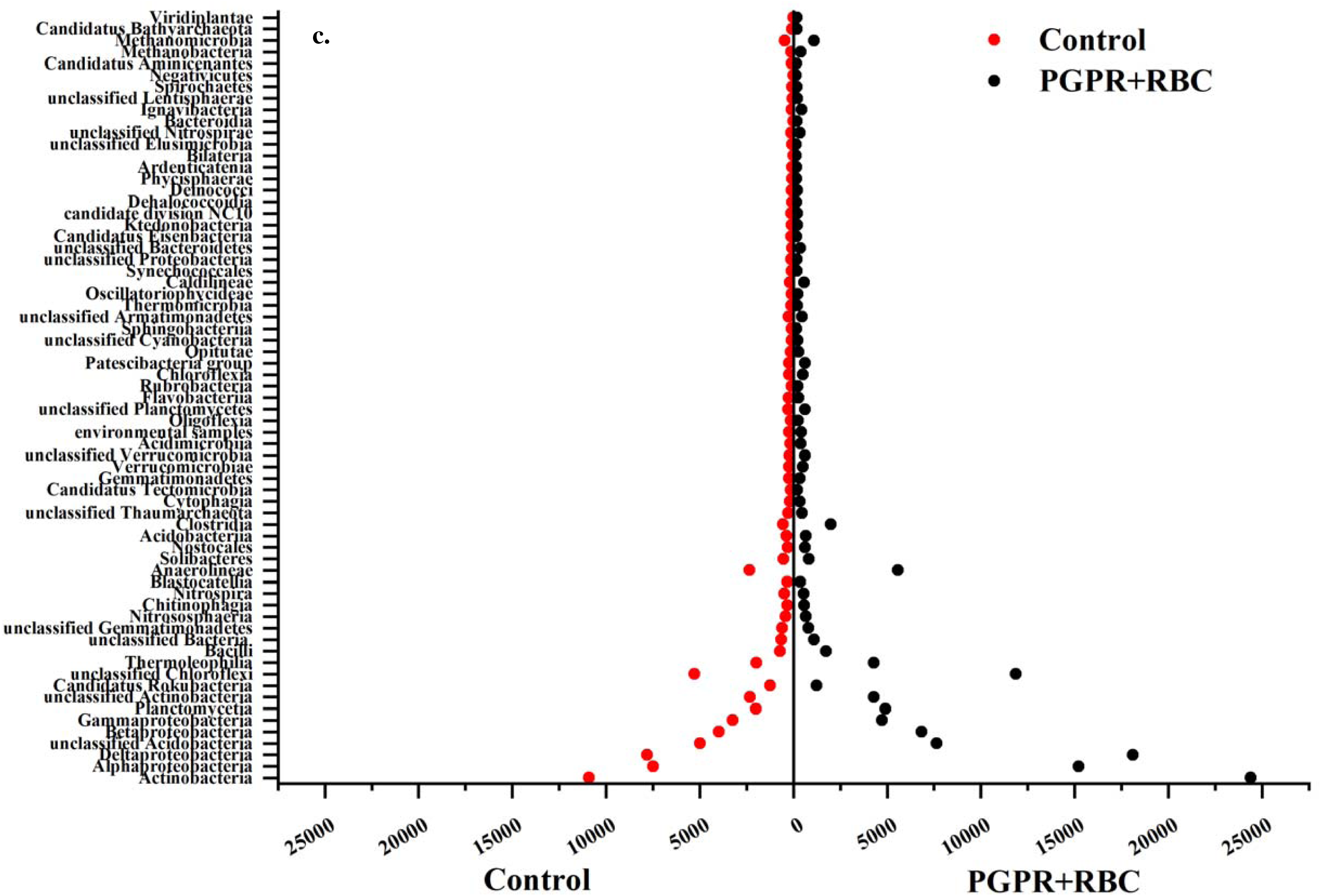
Population pyramid plot of microbial communities under treatments (a-c) compared to the control field microbial communities.

**Table S1.** Instrument specifications during ICP-MS analysis.

| **PARAMETER** | **ICPMS (Parkin-Elmer)** |
| --- | --- |
| **RF Power** | 1.2 Kw |
| **Carrier Gas Flow Rate** | 18 ml/min |
| **Argon Plasma Flow Rate**<br>**(Nebulizer cold Plasma)** | 0.94 L/min |
| **Sampling Depth** | 150 to 180 |
| **Collision Cell Gas** | He (92%) + H (8%) |
| **Collision Cell Gas Flow Rate** | 4-10 ml /min |
| **Nebulizer Pump Rate** | 0.5 rps (30 RPM) |
| **Uptake Time** | 35 s |
| **Wash Time** | 60- 70 s |

**Table S2.** WDXRF general instrumental specifications.

| **Parameters** | **WD XRF (Bruker S8 Tiger)** |
| --- | --- |
| **Analysis range** | Beryllium to Uranium |
| **Sample form** | Powder solid |
| **Concentration range** | Concentrations from sub ppm to 100% |
| **Sample size** | Less than 200mm mess size |
| **Excitation** | End window 4kW Rh anode tube (60Kv, 170mA) |
| **Detector gas** | P10 gas (10 % methane, 90 % argon) |
| **X-ray Detectors** | Gas flow proportional counters and Scintillation counters |

**Table S3.** The procedure of sequential extraction fractionation for assessment of bioavailability.

| Fraction | Solution | Applied Conditions |
| --- | --- | --- |
| F1: Exchangeable | 8 ml of $\text{MgCl}_2$ (1M, pH=7.0) | 1 hr. at Room temperature, continuous agitation |
| F2: Carbonates | 8ml of NaOAc (1M, pH=5.0) w/ $\text{CH}_3\text{COOH}$ | 5 hrs. at Room temperature, continuous agitation |
| F3: Iron and Manganese Oxides | 20ml of $\text{NH}_2\text{OH}\cdot\text{HCl}$ in 25% (v/v) HOAc | 6 hrs. at 96°C, intermittent agitation |
| F4: Organic Matter | 3ml of $\text{HNO}_3$ (0.02M) | 2 hrs. at 85°C, intermittent agitation |
| | 5ml of 30% $\text{H}_2\text{O}_2$ (pH=2) w/$\text{HNO}_3$ | |
| | 3ml of 30% $\text{H}_2\text{O}_2$ (pH=2) w/$\text{HNO}_3$ | 3hrs. at 85°C, intermittent agitation |
| | 5ml of $\text{NH}_4\text{OAc}$(3.2M) in 20% (v/v)$\text{HNO}_3$ diluted to 20mL with Milli-Q water | 30 mins. at Room temperature, continuous agitation |
| F5: Residual | HF-$\text{HClO}_4$ (5 : 1) | - |

**Table S4.**
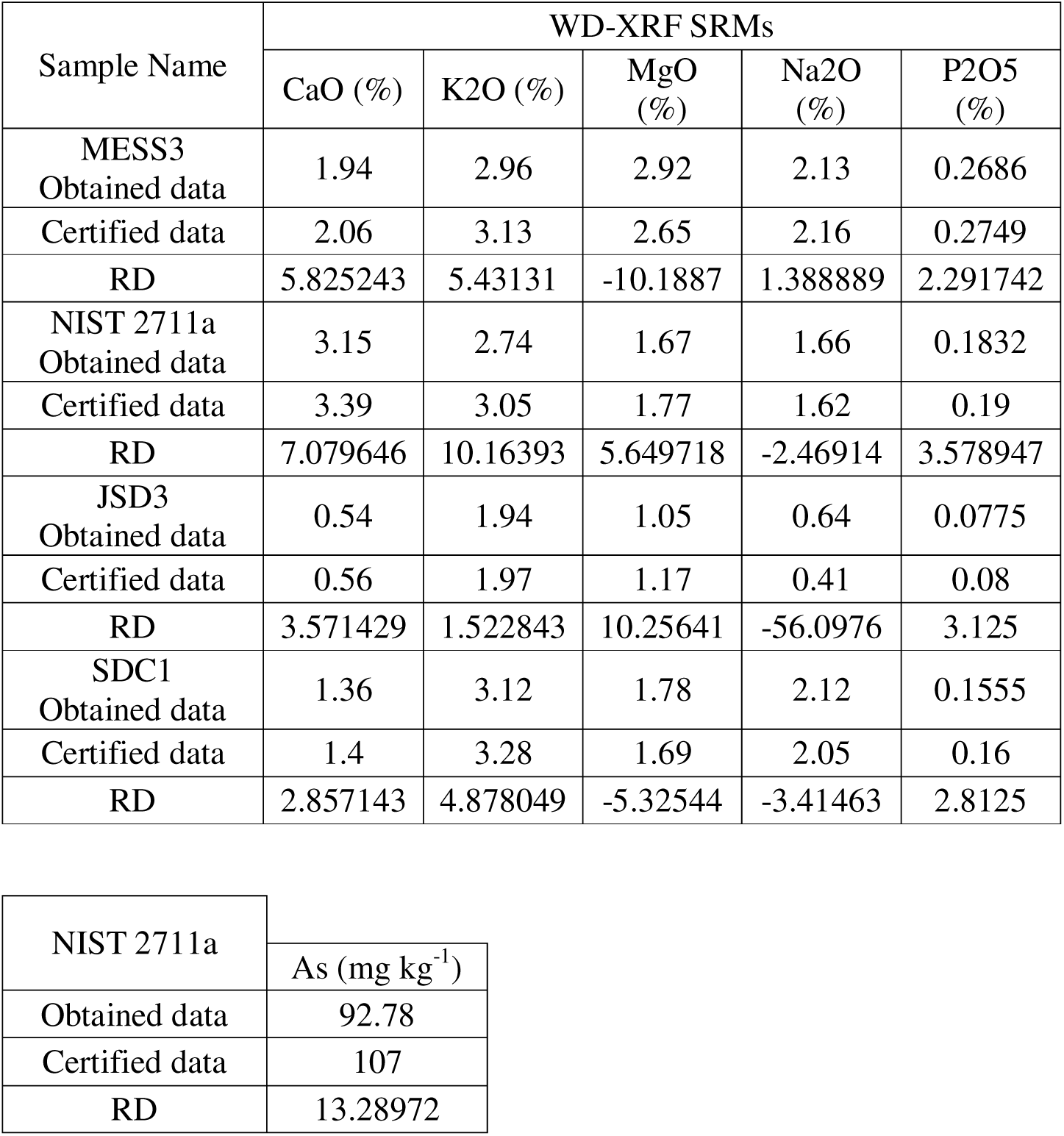
Standard reference material values as analyzed in ICP-MS and WD-XRF.

| Sample Name | WD-XRF SRMs |  |  |  |  |
| --- | --- | --- | --- | --- | --- |
|  | CaO (%) | K<sub>2</sub>O (%) | MgO (%) | Na<sub>2</sub>O (%) | P<sub>2</sub>O<sub>5</sub> (%) |
| MESS3<br>Obtained data | 1.94 | 2.96 | 2.92 | 2.13 | 0.2686 |
| Certified data | 2.06 | 3.13 | 2.65 | 2.16 | 0.2749 |
| RD | 5.825243 | 5.43131 | -10.1887 | 1.388889 | 2.291742 |
| NIST 2711a<br>Obtained data | 3.15 | 2.74 | 1.67 | 1.66 | 0.1832 |
| Certified data | 3.39 | 3.05 | 1.77 | 1.62 | 0.19 |
| RD | 7.079646 | 10.16393 | 5.649718 | -2.46914 | 3.578947 |
| JSD3<br>Obtained data | 0.54 | 1.94 | 1.05 | 0.64 | 0.0775 |
| Certified data | 0.56 | 1.97 | 1.17 | 0.41 | 0.08 |
| RD | 3.571429 | 1.522843 | 10.25641 | -56.0976 | 3.125 |
| SDC1<br>Obtained data | 1.36 | 3.12 | 1.78 | 2.12 | 0.1555 |
| Certified data | 1.4 | 3.28 | 1.69 | 2.05 | 0.16 |
| RD | 2.857143 | 4.878049 | -5.32544 | -3.41463 | 2.8125 |

|  |  |
| --- | --- |
| NIST 2711a | As (mg kg<sup>-1</sup>) |
| Obtained data | 92.78 |
| Certified data | 107 |
| RD | 13.28972 |

**Table S5.**
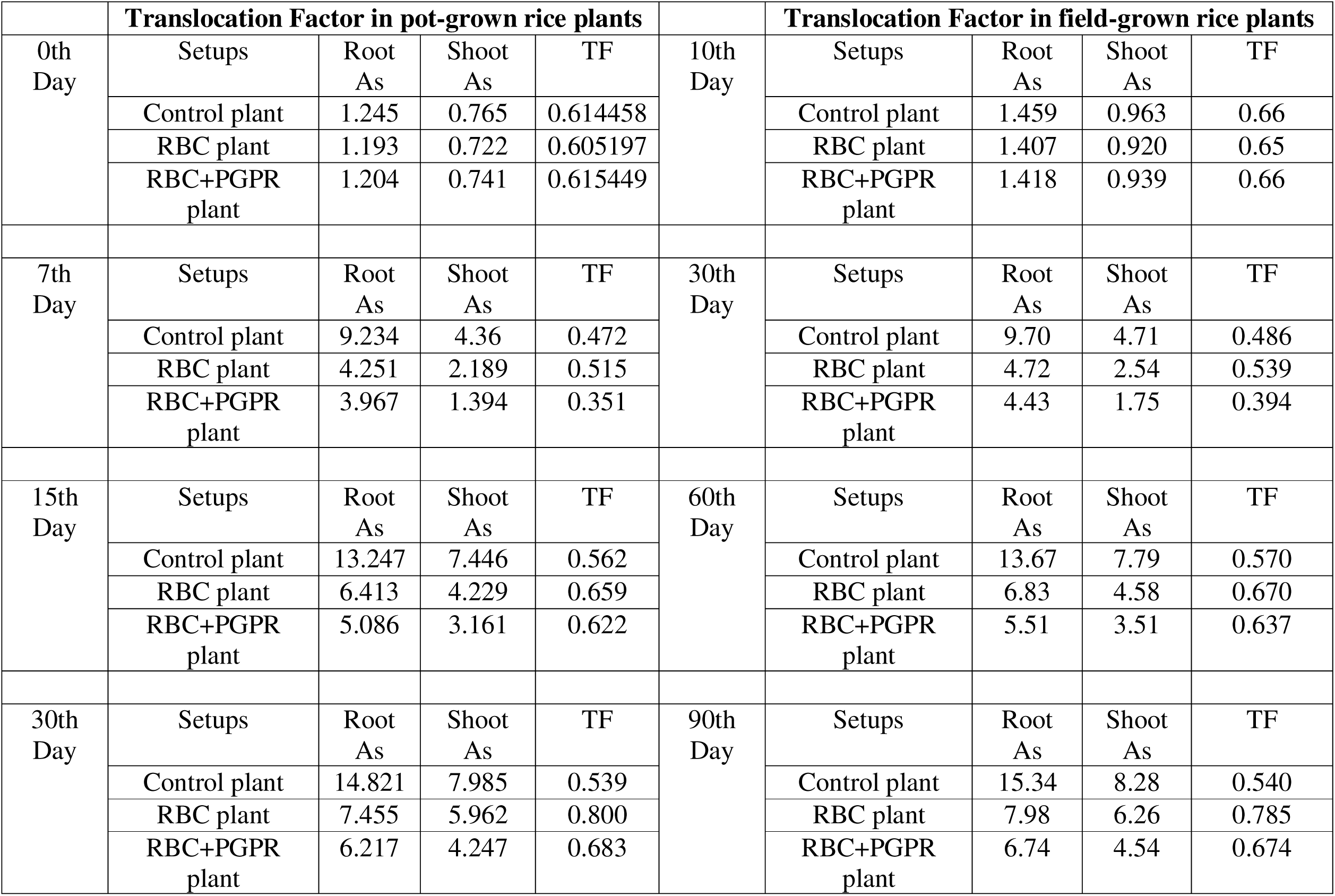
Translocation factors of As for pot-grown and field-grown rice plants under control and treatment setups.

## Notes

### Competing Interest Statement

The authors have declared no competing interest.

